# Measles vaccine attenuation comprises innate immune cell infection and interferon responses

**DOI:** 10.1101/2025.06.09.657851

**Authors:** Felix G.M. Andres, Sebastian Parusel, Oliver Siering, Francisco M. Acosta, Dominic A. Skeele, Katayoun Ayasoufi, Roberto Cattaneo, Bevan Sawatsky, Christian K. Pfaller

## Abstract

Measles virus (MeV) is a highly contagious pathogen capable of infecting immune cells and thereby causing immunosuppression and immune amnesia. A live-attenuated MeV vaccine is highly efficacious and has been safely administered for over 60 years. However, a detailed mechanistic understanding of the mode of its attenuation remains elusive. Here, we determined the molecular mechanisms underlying MeV vaccine attenuation in primary human immune cells. Enhanced infection of innate immune cells resulting in uncontrolled viral RNA synthesis and reduced ability to evade host antiviral responses induced a strong type-I interferon signature that arrested replication of the vaccine strain. In contrast, the wild type virus replicated slowly but steadily, avoiding detection by innate immunity.

## Introduction

Measles virus (MeV) of the *Morbillivirus* genus in the *Paramyxoviridae* family, the causative agent of measles, is the most contagious human pathogen known to date ^1^. The measles basic reproduction number (R_0_), which describes the number of successful transmissions from an infected person to contact persons, is between 12 and 18 ^1,2^. Despite being classified as a respiratory virus, the MeV life cycle is bi-phasic: MeV initially infects immune cells expressing the entry receptor CD150 (or signaling lymphocyte activation molecule, SLAM) ^3^. After extensive replication in CD150-positive B lymphocytes, T lymphocytes, and myeloid cells within primary and secondary immune organs ^4^, the virus is transported back to the airway epithelium where it infects nectin-4-positive cells from the basolateral side ^5,6^.

Large infectious centers form during multiple rounds of replication in these cells ^7^, and dislodged cell clusters containing large amounts of infectious particles may transmit infection to the next host ^8^. Measles is associated with acute immunosuppression ^9,10^ and long-term immune amnesia ^11–13^. Both are directly related to its immune cell-infecting stage resulting in cytokine imbalance ^14–16^ as well as transient leukopenia ^1,9^. Immunosuppression enables opportunistic bacterial secondary infections, which are the main cause of measles-related fatalities ^17^. Long-term loss of memory B cell clones and potentially plasma cell clones is the underlying cause of immune amnesia, which describes the permanent loss of pre-existing antibody responses during the course of an infection with MeV ^12,13^.

Measles is a vaccine-preventable disease ^1,17^. A live-attenuated vaccine was developed by John Enders in the 1950s by passaging a wild type (WT) isolate on mammalian cell lines, chicken embryonic fibroblasts, and in embryonated chicken eggs ^18^; it was approved in 1963 ^19,20^. Today, it is provided as a combination vaccine against measles, mumps, and rubella (MMR vaccine) ^21^. It is 93% effective after a single dose and 97% effective after two doses ^22,23^ and has a favorable safety profile ^24^. Since introduction of the vaccine, measles cases and measles-related fatalities have significantly dropped worldwide ^25^. It is estimated that the measles vaccination program has prevented over 60 million deaths, mostly of children ^25^. However, measles eradication goals by the World Health Organization (WHO) and the Centers for Disease Control and Prevention (CDC) have not been met due to increasing vaccine hesitancy fueled by misinformation such as falsely causal linkage of the measles vaccine with autism ^26,27^.

Despite being in use for over 60 years now, understanding of the attenuation mechanism is incomplete ^28^, which may contribute to vaccine hesitancy ^27^. It is well-established that mutations in the hemagglutinin (H) gene of the vaccine strain (Tables S1, S2) result in additional recognition of CD46 as a cell entry receptor ^29^. In addition, vaccine MeV induces stronger type-I interferon (IFN)-mediated antiviral responses than WT ^28^. This suggests that mutations in the P/V/C gene products, phosphoprotein (P), V protein, and C protein (Tables S1, S2) may reduce their ability to counteract innate immune responses ^10,30^. To which extent these mutations contribute to the vaccine virus attenuation is not known.

Here, we systematically compared the ability of WT and vaccine MeVs to infect and replicate in primary peripheral blood mononuclear cells (PBMCs) from human donors. We characterized differential host responses underlying the attenuation of vaccine MeV. By generating recombinant viruses in which P/V/C and/or H genes were replaced with their WT counterparts, we show that both genes contribute to the attenuated phenotype of the MeV vaccine. While vaccine H enhances infectivity and tropism in cells from the myeloid compartment, vaccine P/V/C gene products cannot efficiently control induction of type-I IFN-mediated antiviral immunity.

## Results

### Enhanced infection rate in non-B/T/NK cells by vaccine strain MeV correlates with reduced productive infection

To assess the ability of WT and vaccine MeVs to replicate in primary cells, we relied on recombinant viruses IC323 (WT strain) ^31^ and vac2 (vaccine strain) ^32^ expressing a green fluorescent protein (GFP) reporter from an additional transcription unit located between the H and L genes (Fig. 1A). In addition, we included an IC323-derived C protein-deficient (C^KO^) virus previously shown to be attenuated through activation of double-stranded RNA (dsRNA)-mediated innate immune responses ^33,34^. In primary human PBMCs, all viruses replicated to similar titers for 48 h after infection (Fig. 1B). However, only WT virus established productive replication with increasing titers at later time points, whereas titers of both C^KO^ and vac2 did not significantly increase over time (Fig. 1B).

**Fig. 1.**
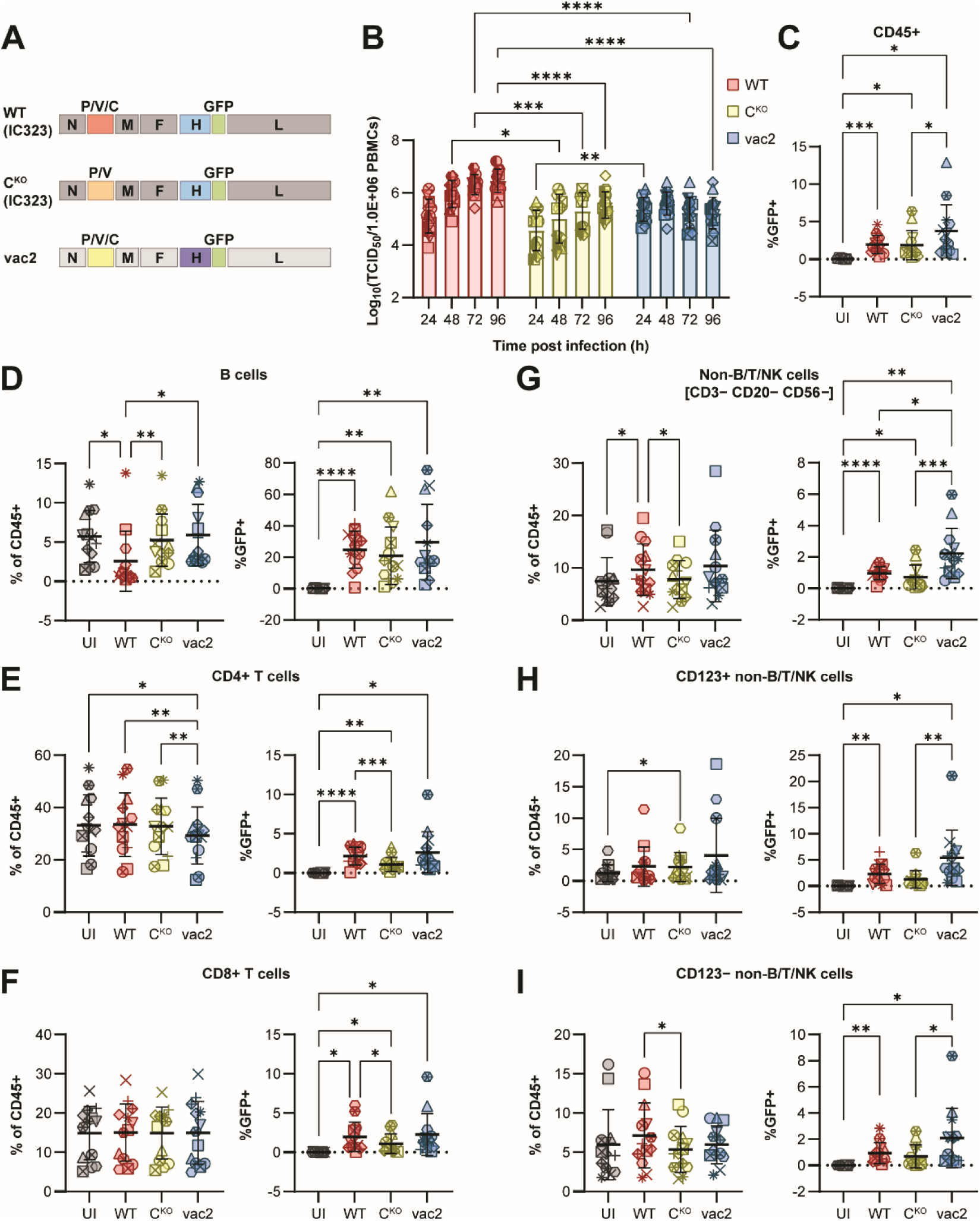
Measles vaccine virus exhibits enhanced infectivity in myeloid cells but stagnating production of infectious progeny. (A) Genome schemata of recombinant viruses used in this study. Wild type (WT) and C protein-deficient (C^KO^) viruses are derived from IC323 strain; vaccine virus (vac2) is derived from Moraten vaccine strain. All viruses express a GFP reporter from an additional transcription unit between H and L genes. (B) Virus growth kinetics in primary PBMCs from human donors (n = 18) infected at multiplicity of infection of 0.1. Statistics: Two-way ANOVA with Tukey’s multiple comparisons test. (C) – (I) Flow cytometry analysis of infection rates based on GFP-reporter expression in different immune cell subsets from human donors (n = 13). Statistics: Repeated measures one-way ANOVA with Tukey’s multiple comparisons test. (C) GFP+ cells of CD45+ cells. (D)-(I) Left diagrams show cell type frequencies in CD45+ population; right diagrams show GFP+ cells of cell type. (D) B cells. (E) CD4+ T cells. (F) CD8+ T cells. (G) Non-B/T/NK (CD3− CD20− CD56−) cells. (H) CD123+ non-B/T/NK cells. (I) CD123− non-B/T/NK cells. In all graphs, each donor is identified with a unique symbol. Bars indicate mean values and error bars show standard deviations. Asterisks indicate statistical significance levels: * *P*≤0.05; ** *P*≤0.01; *** *P*≤0.001; **** *P*≤0.0001.

We next sought to quantify the infection rates of the three viruses in different immune cell subsets. For this, we established a 24-color spectral flow cytometry panel to phenotype different lymphoid and myeloid immune cell subsets (Fig. S1). At 48 h post infection, the time point when virus kinetics indicated substantial difference between WT and vac2 growth (Fig. 1B), we observed similar infection rates in total CD45+ immune cells, although vac2 exhibited a trend for higher infection rates than WT or C^KO^ (Fig. 1C). As expected, infection rates were highest in CD20+ B cells (Fig. 1D), CD4+ T cells (Fig. 1E), and CD8+ T cells (Fig. 1F). While C^KO^ exhibited significantly reduced infection rates in T cells compared to WT, vac2 infected these cells with a tendentially higher rate, independent of their memory status (Fig. S2). Similar trends were observed in γδ-T cells (CD3+ TCRγδ+, Fig. S3A), NKT cells (CD3+ CD56+, Fig. S3B), natural killer (NK) cells (CD20− CD3− CD56+, Fig. S3C), and NK-suppressor cells (CD20− CD3− CD56hi, Fig. S3D), although NK and NK-suppressor cells were most resistant to MeV infection in general. Notably, only WT infection led to significantly lower B cell numbers (Fig. 1D, left diagram), indicative for its ability to deplete B cells. This depletion affected IgD−IgM+ B cells (Fig. S3E), but not other B cell subsets (Fig. S3F-H). Importantly, vac2 exhibited significantly higher infection rates in lineage-negative (CD20− CD3− CD56−) non-B/T/NK cells compared to both WT and C^KO^ (Fig. 1G). This was confirmed across different non-B/T/NK cell subsets (Fig. 1H-I, Fig. S4, Fig. S5).

Finally, we asked whether the relative frequencies of different cell types in individual donor samples had an impact on overall WT or vac2 infections (Fig. S6). For both WT and vac2, higher frequencies of CD4+ T cells (Fig. S6A), and B cells (Fig. S6D) had a positive effect on the overall infection rates in CD45+ cells. In contrast, donors with high NK and NK-suppressor cell frequencies exhibited lower overall infection rates (Fig. S6I, J), which correlated with the generally low infection rates of these cell types (Fig. S3C, D). Notably, γδ-T cells (Fig. S6C), and CD123− non-B/T/NK cells (Fig. S6G) exhibited a positive trend for vac2 but not for WT. Other cell types had no or negative correlation with infectivity of both viruses. These results indicate that vac2 infects more cell types than WT, especially non-B/T/NK cells that are minimally infected by WT.

### MeV vaccine strain induces robust type-I IFN signature and CD8 T cell response

We next asked whether arrest of vac2 replication was associated with induction of antiviral host responses. We performed total bulk RNA-seq of matched samples from three donors infected with WT, C^KO^, or vac2, and determined differential gene expression compared to uninfected cells (UI). Principal component analysis revealed strong donor effects, but nevertheless, vac2 samples consistently deviated more strongly from UI samples than WT or C^KO^ samples (Fig. 2A), and only vac2 infection outweighed donor-to-donor variations in the hierarchical clustering (Fig. 2A, B). While vac2 infection induced a consistent differential gene expression pattern across the donors, WT and C^KO^ infections induced only mild changes in gene expression patterns (Fig. 2B). Overall, WT infection led to significant downregulation of only four genes without significantly upregulating genes (Fig. 2C). In contrast, C^KO^ infection upregulated 69 genes and downregulated 40 genes significantly. Strikingly, vac2 infection had a powerful impact on the cellular transcriptome inducing significant upregulation of 1385 and downregulation of 1141 genes (Fig. 2C). Among the top 20 genes upregulated during C^KO^ infection were immune-related genes such as *CCL18*, *DCSTAMP*, *ALDH1A2*, and *CD38* (Fig. 2D). For infection with vac2, however, we identified *IFNB1* and *IFNL2* genes as top upregulated genes, and the IFN response-related genes such as *ZBP1* and *ISG20* were among the top 20 upregulated genes (Fig. 2E). Most of the top genes upregulated during vac2 infection were not or only mildly upregulated during WT or C^KO^ infection (Fig. 2E, lower diagram). Among top downregulated genes, we found that interleukin-17 family members *IL17A* and *IL17F* were significantly downregulated by all three viruses (Fig. S7A-C).

**Fig. 2.**
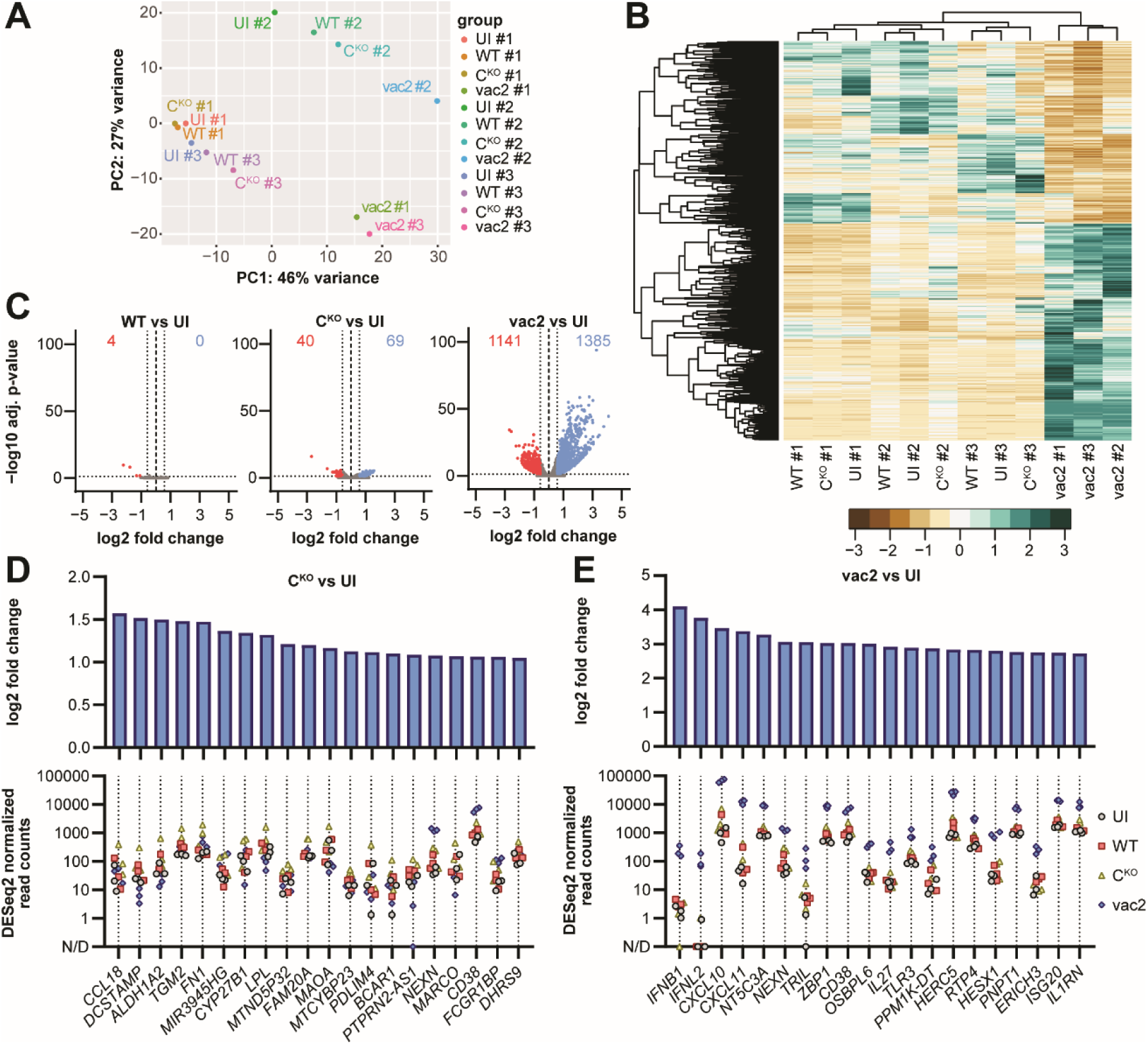
Measles vaccine induces strong differential gene expression in PBMCs. RNA-seq was performed with 3 donors and 4 conditions per donor: uninfected (UI), WT infected, C^KO^ infected, vac2 infected. (A) Principal component analysis of normalized gene count tables calculated by DESeq2. (B) Z-score plot of differentially expressed genes. (C) Volcano plots summarizing adjusted p-values and fold changes of differentially expressed genes of virus-infected versus UI. Red: significantly downregulated; blue: significantly upregulated; grey: not significant. Numbers indicate the number of genes in the respective quadrant. (D) Top 20 upregulated genes during C^KO^ infection, ranked by decreasing log2(fold change) (top diagram). The bottom diagram indicates the normalized read counts of each individual sample. (E) Top 20 upregulated genes during vac2 infection, ranked by decreasing log2(fold change) (top diagram). The bottom diagram indicates the normalized read counts of each individual sample.

To confirm that vac2 infection induced a strong type-I IFN signature in PBMCs, we performed gene ontology (GO) analysis on both significantly upregulated (Fig. 3A, B) and downregulated genes (Fig. S7D-F). GO terms strongly associated with upregulated genes during vac2 infection included “response to virus”, “response to interferon-beta”, and similar terms (Fig. 3A). GO terms associated with upregulated genes during C^KO^ infection included “innate immune response”, and “cellular response to cytokines”, indicating that both viral infections induced immune responses, but to different degrees. No strong association with specific GO terms was found with downregulated genes in response to either WT, C^KO^, or vac2 infection (Fig. S7D-F).

**Fig. 3.**
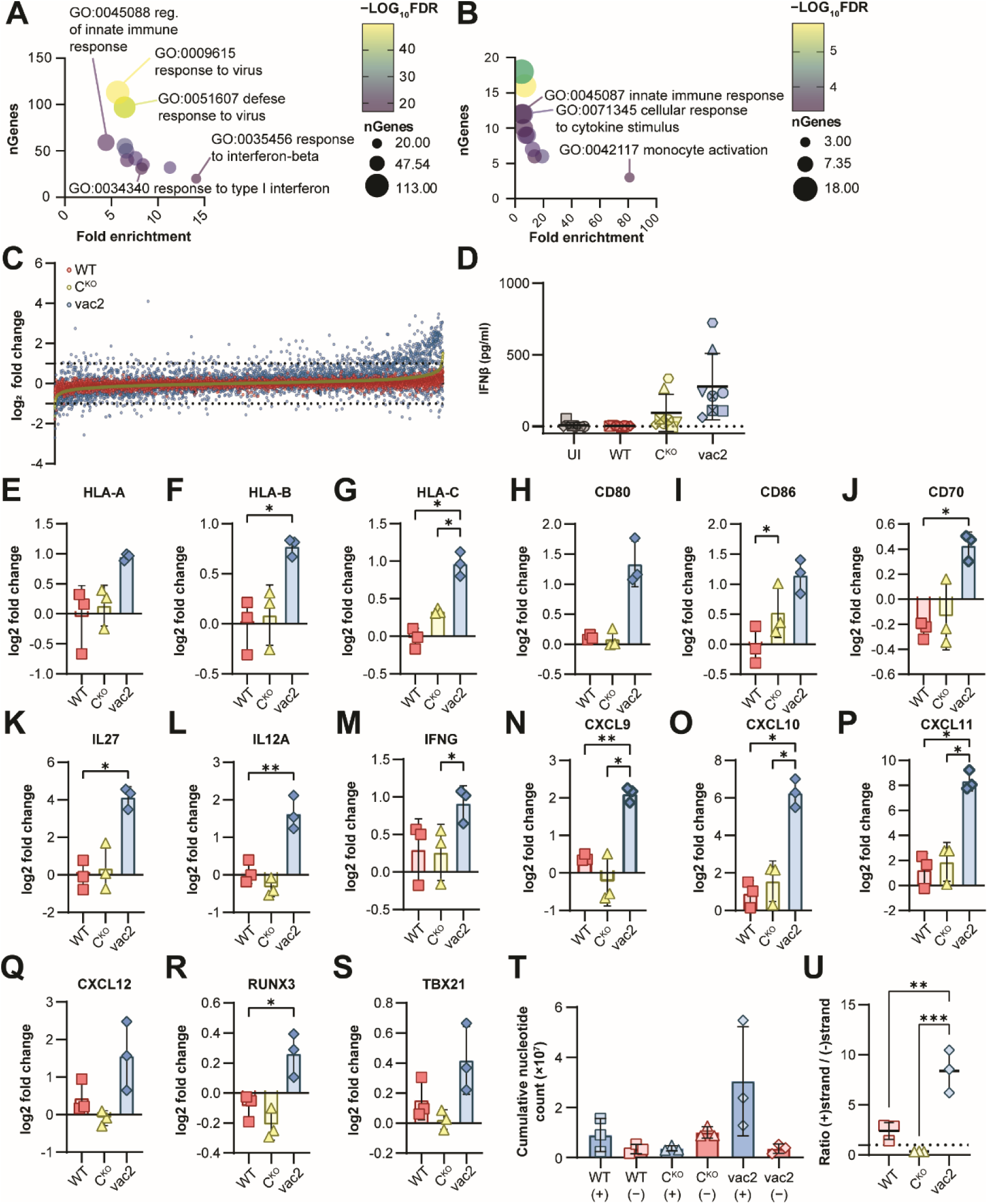
Measles vaccine strain induces strong type-I interferon response. (A)-(B) GO term analysis of genes upregulated during infection. Circle color indicates −log10(FDR) determined by ShinyGO. Circle size is proportional to the number of genes (nGenes) identified by ShinyGO. (A) vac2 infection. (B) C^KO^ infection. (C) Log2(fold change) of ISG expression after infection with different viruses. ISGs are ranked along x-axis from left to right in ascending order for C^KO^ infection (yellow), a known inducer of type-I IFN responses in cell lines. Dotted lines indicate 2-fold up-or downregulation. (D) Human IFNβ ELISA of supernatants from PBMCs collected 48 h post infection (n = 8). Bars indicate mean values and error bars show standard deviations. Statistics: Repeated measures one-way ANOVA with Tukey’s multiple comparisons test. (E)-(S) Log2 fold change of mRNA levels of virus-infected samples compared to uninfected derived from RNA-seq data for genes involved in T cell-responses. (E)-(G) MHC-I. (H)-(J) Co-stimulatory molecules. (K)-(M) Signal 3 cytokines. (N)-(Q) Effector chemokines. (R)-(S) IFNγ-activated genes. (T) Cumulative nucleotide counts of reads aligning to (+)strand or (−)strand viral genomic sequence. (U) Ratio of (+)strand-aligning over (−)strand-aligning nucleotide counts. The dotted line is at 1.Statistics: one-way ANOVA [repeated measures in panels (E)-(S)] with Tuckey’s multiple comparisons test. Bars indicate mean values and error bars show standard deviations. Asterisks indicate statistical significance levels: * *P*≤0.05; ** *P*≤0.01; *** *P*≤0.001.

Finally, we utilized the Interferome Database ^35^ to identify 6786 type-I IFN-stimulated genes among 33720 total genes from our RNA-seq analyses. 654 ISGs were upregulated at least 2-fold during vac2 infection, and 31 during C^KO^ infection (Fig. 3C). IFN-beta (IFNβ) ELISA of supernatants harvested at 48 h post infection supported our finding that C^KO^ induced a moderate IFN response whereas vac2 induced a robust IFN response (Fig 3D). Notably, WT infection did not induce any secretion of IFNβ, supporting our finding that the antiviral defense system of PBMCs is blind to this virus.

Besides a strong type-I IFN signature, vac2 infection induced signals required for CD8 T cell responses on a transcriptional level: MHC-I gene expression (Fig. 3E-G), co-stimulatory molecules *CD80*, *CD86*, and *CD70* (Fig. 3H-J), signal 3 cytokines *IL27*, *IL12A*, and *IFNG* (Fig.3K-M), effector chemokines *CXCL9*, *CXCL10*, *CXCL11*, and *CXCL12* (Fig. 3N-Q), and IFNγ signaling-related genes *RUNX3* and *TBX21* (Fig. 3R-S) were all upregulated. In contrast, MHC-II genes were not upregulated but rather trended to virus-induced downregulation (Fig. S7G-O). Expression of the co-stimulatory molecule *CD40* and cytokines *IL12B* and *IL2* remained unchanged (Fig. S7P-R). These data indicate that vac2 infection, but not WT-strain infection, induces CD8 T cell responses. Notably, C^KO^ infection also did not induce CD8 T cell responses to the extent of vac2 infection.

In addition to host gene expression, we quantified viral RNA to determine whether the transcription of the three viruses differed. (+)strand (mRNA, antigenome) coverage of vac2 was higher compared to WT, whereas C^KO^ exhibited lower coverage (Fig. 3T, Fig. S8A).

Interestingly, the (−)strand (genome) coverage of C^KO^ appeared slightly elevated compared to WT (Fig. 3T, Fig. S8A). While WT virus (+)strand reads were more abundant than (−)strand reads, this ratio was reversed for C^KO^ (Fig. 3U), suggesting that C^KO^ may have transcriptional defects in primary PBMCs. Interestingly, the transcriptional gradient of viral mRNA species was flattened for C^KO^, but not for WT or vac2 (Fig. S8B). Importantly, vac2 had significantly higher ratio of transcripts to genomes compared to WT (Fig. 3U), suggesting that this virus initially transcribes efficiently but eventually fails to produce negative strand genome and complete replication.

### Wildtype P/V/C and H genes affect vaccine strain replication differently

We next sought to determine the genetic etiology underlying the differential host responses against WT and vac2 viruses. We thought that H would mediate the enhanced infection rate of vac2 in non-B/T/NK cells whereas P, V, and C proteins would impact immune evasion. Amino acid mutations occurring in each of these proteins are partially associated with functional differences (Table S2). We generated three vac2-based chimeric viruses in which we replaced the vac2 P/V/C gene with its WT analog (vPwt), the vac2 H gene (vHwt), or both (vPHwt; Fig. 4A). Growth kinetics analyses revealed that vPwt replication was slightly enhanced compared to vac2, despite one donor that supported vac2 replication extraordinarily well (Fig. 4B, upside triangle). However, vPwt did not reach the titers of WT (Fig. 4B). In contrast, vHwt replication was more limited than that of vac2 (Fig. 4B), indicating that H protein of WT indeed acts as a restriction factor limiting viral entry into CD150+ cells, whereas H protein of vac2 extends cell tropism to CD150− CD46+ cells. Notably, WT P/V/C enhanced replication also in combination with WT H (Fig. 4B), indicating that the two gene functions were independent of each other.

**Fig. 4.**
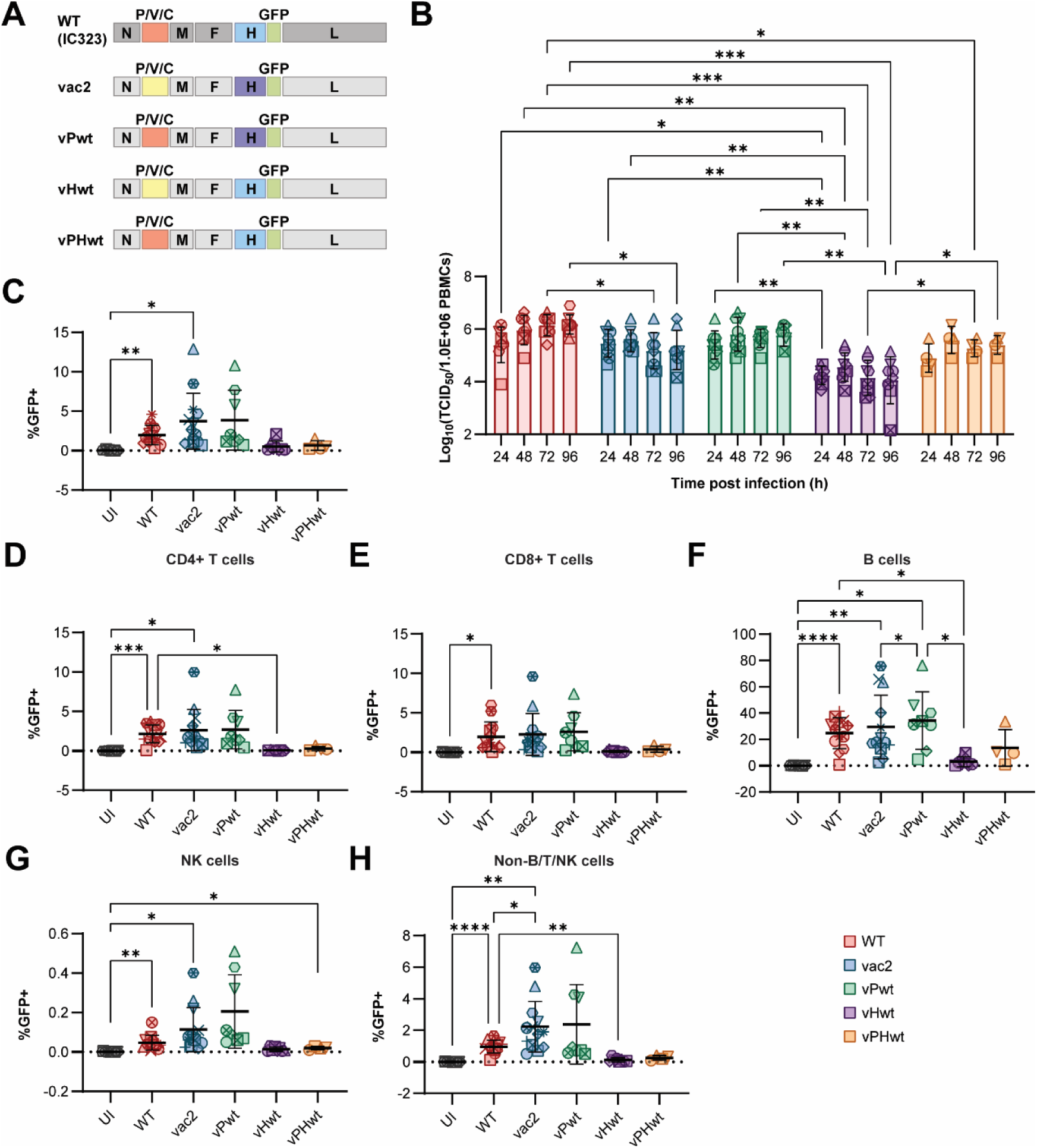
Measles P/V/C and H gene products affect the ability to productively replicate in PBMCs. (A) Schematic representation of chimeric vaccine viruses expressing WT P/V/C (vPwt), WT H (vHwt), or both WT P/V/C and H (vPHwt). (B) Virus growth kinetics in primary PBMCs from human donors (n = 8, except vPHwt: n = 4) infected at multiplicity of infection of 0.1. Statistics: Two-way ANOVA with Tukey’s multiple comparisons test. (C) – (H) Flow cytometry analysis of infection rates based on GFP-reporter expression in different immune cell subsets from human donors 48 h post infection (WT, vac2: n = 13; vPwt, vHwt: n = 8; vPHwt: n = 4). Statistics: Repeated measures one-way ANOVA with Tukey’s multiple comparisons test. (C) GFP+ cells of CD45+ cells. (D) GFP+ cells of CD4+ T cells. (E) GFP+ cells of CD8+ T cells. (F) GFP+ cells of B cells. (G) GFP+ cells of NK cells. (H) GFP+ cells of non-B/T/NK cells. In all graphs, each donor is identified with a unique symbol. Bars indicate mean values and error bars show standard deviations. Asterisks indicate statistical significance levels: * *P*≤0.05; ** *P*≤0.01; *** *P*≤0.001; **** *P*≤0.0001. Note: Data points for UI, WT, and vac2 in (B)-(H) are also included in corresponding diagrams in Fig. 1.

We next determined whether infection rates and cell tropism of the chimeric viruses were altered. vPwt exhibited infection rates in total hematopoietic cells (CD45+ PBMCs) like vac2 and higher than WT (Fig. 4C, Fig. S9A). In contrast, both vHwt and vPHwt infected cells at lower rates than WT (Fig. 4C, Fig. S9A). We observed the same patterns for different cell types (Fig. 4D-H). Notably, total B cell numbers were reduced with vPHwt, as previously observed with WT (Fig. S9B), while cell type frequencies for CD4+ T cells, CD8+ T cells, NK cells, and non-B/T/NK cells were not affected by any viral infection (Fig. S9C-F).

To determine whether the altered replication efficiency of chimeric MeVs was related to their ability to evade innate immune responses, we examined IFNβ secretion at 48 h p.i. (Fig. 5A). Indeed, vPwt-infected PBMCs secreted reduced levels of IFNβ into supernatants than vac2-infected cells, and vPHwt infection resulted in lower IFNβ levels than vHwt infection.

**Fig. 5.**
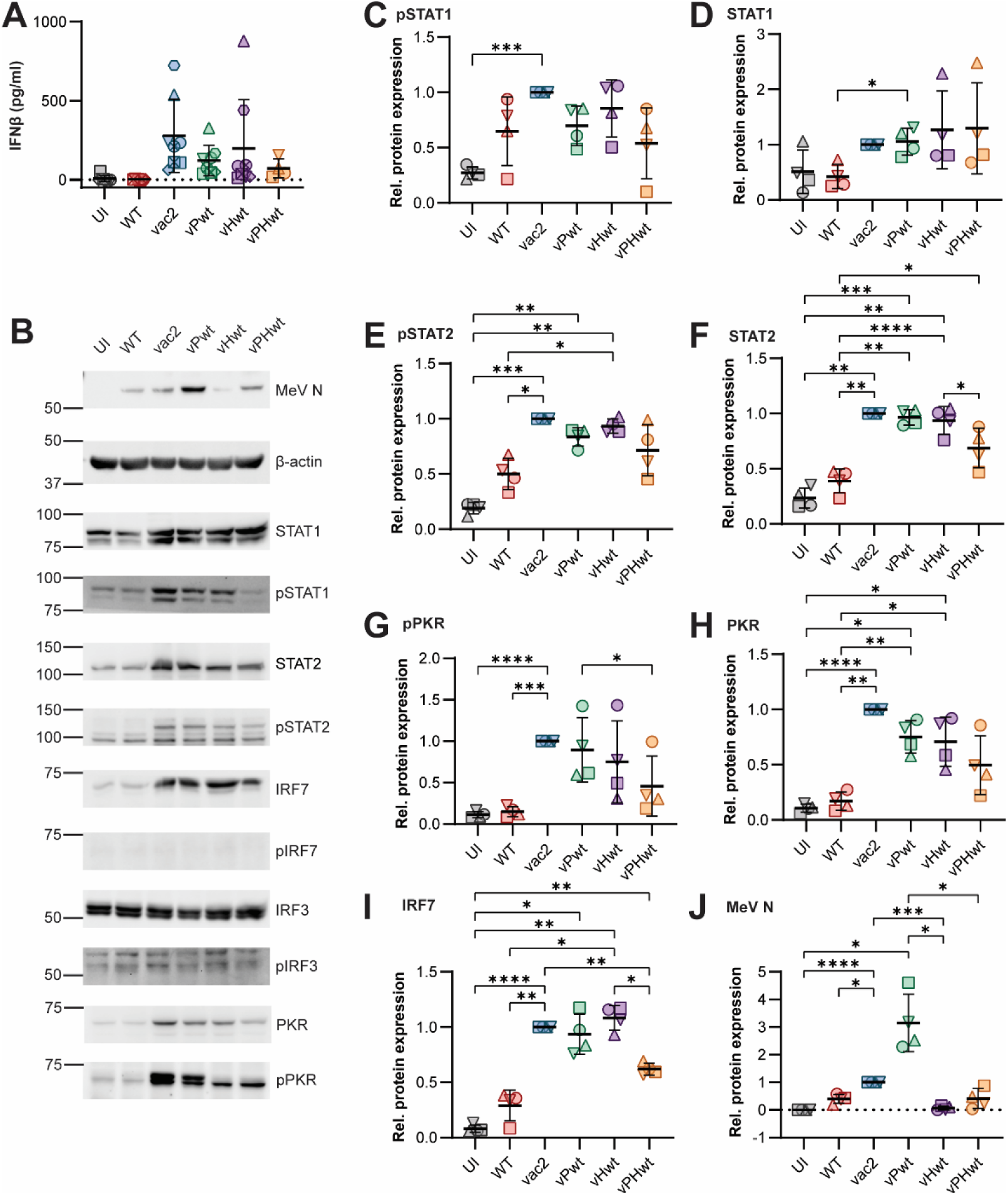
Measles vaccine backbone activates innate antiviral responses. (A) Human IFNβ ELISA of supernatants from PBMCs collected 48 h post infection (UI, WT, vac2, vPwt, vHwt: n = 8; vPHwt: n = 4). Bars indicate mean values and error bars show standard deviations. Statistics: Repeated measures one-way mixed effects model with Tukey’s multiple comparisons test. Note: Data points for UI, WT, and vac2 are also included in corresponding diagrams in Fig. 1. (B) Western blot analysis of viral protein expression (N protein) and innate immunity activation in PBMCs 48 h post infection with indicated viruses. (C)-(J) Quantifications of western blots shown in (B) and Fig. S10. Bars indicate mean values and error bars are standard deviations. Statistics: One-way ANOVA with multiple comparisons test. (C) phospho-STAT1 (pSTAT1). (D) Total STAT1. (E) phospho-STAT2 (pSTAT2). (F) Total STAT2. (G) phospho-PKR (pPKR). (H) Total PKR. (I) Total IRF7. (J) MeV N protein. Statistics: Repeated measures one-way mixed effects model with Tukey’s multiple comparisons test. Asterisks indicate statistical significance levels: * *P*≤0.05; ** *P*≤0.01; *** *P*≤0.001; **** *P*≤0.0001.

However, none of the chimeric viruses completely abolished IFNβ production like WT, indicating that the IFN-antagonistic functions of P, V, and C proteins, while being important factors for this purpose, may not be fully sufficient to perform this action.

To test whether IFN induction or IFN signaling pathways were activated by individual viruses, we performed western blot analyses for activation of transcription factors IRF3, IRF7, STAT1, STAT2, and PKR (Fig. 5B-J, Fig. S10). While phospho-IRF3 (pIRF3_S396_) and phospho-IRF7 (pIRF7) were minimally present at 48 h post infection, phospho-STAT1 (pSTAT1) and phospho-STAT2 (pSTAT2) levels increased in vac2-infected cells compared to WT-infected cells (Fig. 5B, C and E, Fig. S10). Notably, no significant differences were observed between vac2 and the three chimeric viruses. All exhibited enhanced STAT1/STAT2 activation, aligning with upregulation of total STAT1, and STAT2 levels compared to UI or WT-infected cells (Fig. 5B, D, F, Fig. S10). Expression of total PKR and IRF7 was upregulated in vac2-infected cells but not in WT-infected cells (Fig. 5B, H, I, Fig. S10), and so was PKR activation (Fig. 5B, G, Fig. S10). Interestingly, chimeric vac2 viruses, especially vPHwt, exhibited reduced PKR phosphorylation (Fig. 5B, G, Fig. S10) and total PKR upregulation (Fig. 5B, H, Fig. S10). Finally, viral N protein levels were increased in vac2-infected cells compared to WT (Fig. 5B, J, Fig. S10). Chimeric viruses expressing WT P/V/C, especially vPwt, exhibited enhanced N protein levels compared to their respective counterparts (compare vPwt to vac2, and vPHwt to vHwt). Thus, even when two WT genes were introduced in the vaccine genome, this genome retained some attenuation properties.

## Discussion

WT MeV infection is associated with lymphopenia, cytokine imbalance, immunosuppression, and immune amnesia ^10,12,13,36^. None of these clinical signs are observed after MeV vaccination in otherwise healthy individuals ^37^, but molecular insights on the attenuation mechanisms are missing. Here, we show that the MeV vaccine is highly immunogenic in primary PBMCs, inducing strong type-I IFN-mediated antiviral responses. In contrast, we discovered that WT MeV replicates in PBMCs without inducing an innate immune response; WT MeV can control its replication process and avoid accumulation of immunostimulatory RNAs (isRNAs). We have identified three key processes contributing to vaccine attenuation (Fig. 6).

**Fig. 6.**
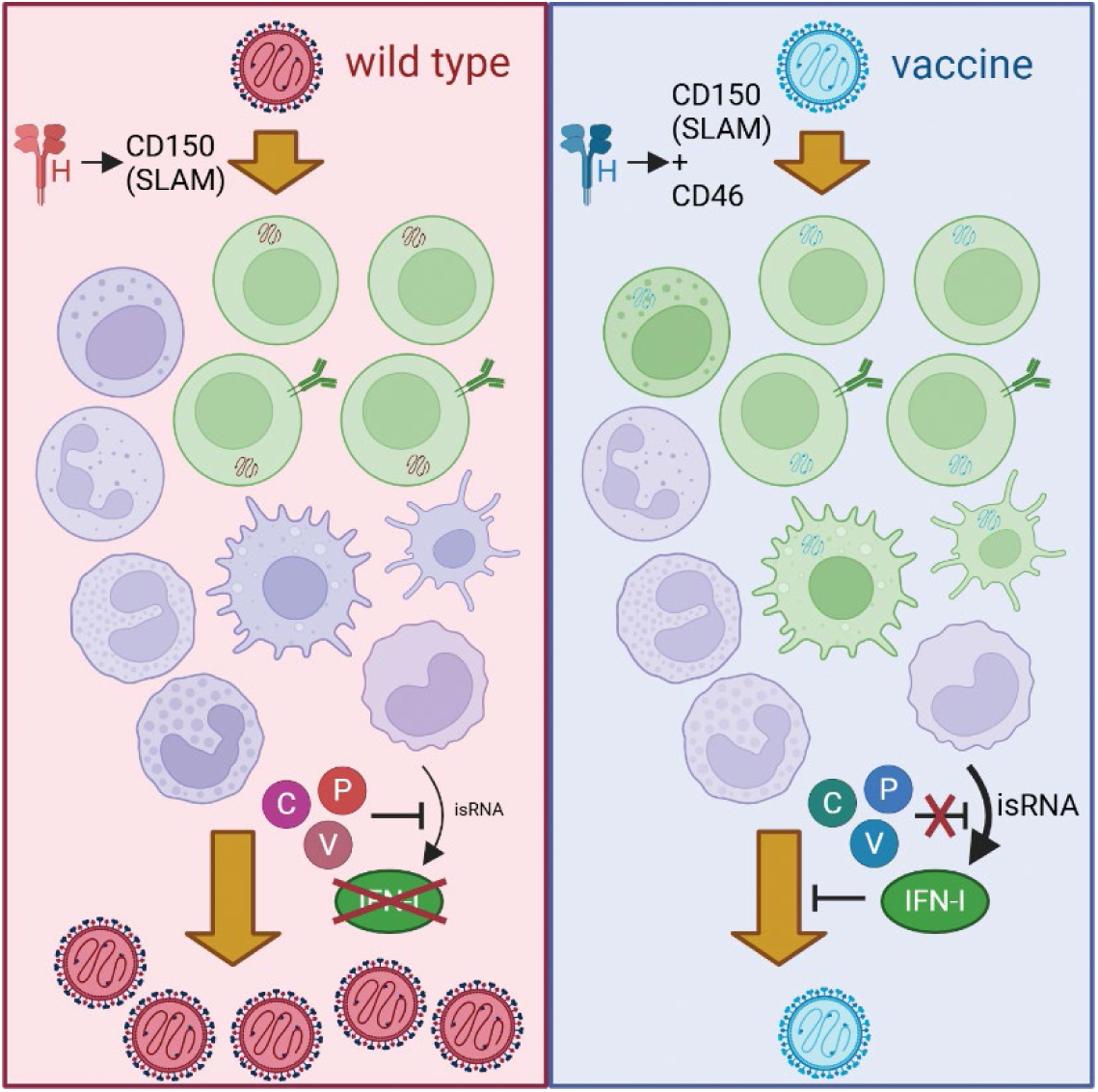
Mechanism of measles vaccine attenuation involves functional changes in H and P/V/C gene products. WT H limits virus entry to CD150+ cells, whereas vaccine H extends tropism to CD46+ cells including some non-B/T/NK cell subsets. WT P/V/C gene products efficiently inhibit type-I IFN responses, whereas vaccine P/V/C gene products are less efficient. Moreover, additional genetic differences between vaccine and WT strains render vaccine more immunogenic, presumably by generating higher levels of immunostimulatory RNA (isRNA). Consequently, WT replication is highly controlled and occurs unnoticed to the innate immune system, whereas vaccine replication is immunogenic and self-limiting. Created in BioRender. Pfaller, C. (2025) https://BioRender.com/g2ei0s4.

First, the MeV vaccine is less discriminating than the WT regarding cell entry. This is caused by mutations in the H gene allowing CD46-dependent cell entry ^29^. We found significantly higher infection rates of vaccine MeV in non-B/T/NK cells compared to WT. This increase was found in both CD123− and CD123+ cells. While our marker panel does not allow exact subtype determination, the CD123+ CD11b− CD11c− cell subset includes plasmacytoid dendritic cells (pDCs), which are major pathogen-sensing innate immune cells ^38,39^.

Second, vaccine MeV replication activates a strong innate antiviral response driven by immediate viral RNA synthesis. In direct comparison, MeV vaccine generated a stronger type-I IFN signature in PBMCs than the C^KO^ mutant of the WT strain. While this was surprising to us, since C^KO^ derivatives of WT or vaccine lineage evoke more robust innate immune responses than their C protein-expressing counterparts in cell lines ^33,34,40,41^, growth kinetics of the two viruses indicate that C^KO^ exhibits a growth defect immediately after entry in primary cells whereas transcription of vaccine MeV starts efficiently but induces a strong antiviral response. We think that preventing innate antiviral responses is a prerequisite for the success of WT MeV given the fact that this virus evolved to replicate within the immune cell compartment for several days before being transmitted by the host. Similarly, Epstein-Barr virus (EBV) does not efficiently induce type-I IFN responses in B cells, which are its primary target, while pDCs respond to EBV infection with production of IFNα ^42^.

The third key to efficient host innate immune evasion are highly functional IFN antagonists. MeV has evolved three proteins with such functions, all encoded in the P/V/C gene^10^. Phosphoprotein (P), which is primarily a polymerase cofactor ^1^, binds and inhibits STAT1 phosphorylation ^43^. V protein blocks multiple innate immune factors including the double stranded RNA (dsRNA) sensor MDA-5 ^44,45^, as well as transcription factors IRF7 ^46^ and STAT2^47^. C protein blocks the transcriptional activity of IRF3 ^48^ and limits the amount of immunostimulatory dsRNA generated during viral RNA synthesis ^33,34^. Our data indicate that the WT-encoded proteins block innate immunity activation more efficiently than their vaccine-encoded counterparts. Specifically, chimeric viruses encoding WT P/V/C led to decreased IFNβ production compared to the analogous viruses encoding vaccine P/V/C. Notably, activation of innate immune markers IRF3, IRF7, STAT1, and STAT2 was not significantly altered between infections with chimeric vaccine strain viruses encoding WT or vaccine P/V/C genes. This suggests that most functions of P and V proteins are preserved in vaccine viruses. This is not surprising, since many of these functions were originally discovered with vaccine strain-derived proteins and key residues involved in immune evasion are conserved between WT and vaccine strains ^43,47,49^. In contrast, increased IFNβ secretion by vaccine MeV is likely the result of loss of the C protein function ^48^. Only WT C efficiently blocks IFNβ transcriptional upregulation, and this requires a functional nuclear localization signal in C protein ^48^. Importantly, expression of WT P/V/C proteins by vaccine MeV did not fully rescue its replication deficiency compared to WT MeV, indicating that P/V/C-mediated immune evasion mechanisms are not sufficient to cancel out the immunostimulatory processes elicited by the vaccine strain. Thus, WT MeV may encode additional functions preventing the accumulation of isRNAs during replication. Notably, the nucleocapsid (N) protein encapsidating viral genomic RNA, and the catalytic subunit of the viral RNA-dependent RNA polymerase (L protein) harbor several additional differences between WT and vaccine strains that may be involved in attenuation (Table S1). Additional studies are required to address their role in vaccine attenuation.

Measles vaccine attenuation involved three key processes performed by different genes. Reversion of individual genes to WT sequences did not fully restore the infectivity of the vaccine strain in PBMCs. This is a critical point speaking for the safety of the vaccine, since reversion to WT, as observed with other live-attenuated vaccines in the past ^50^ would require may, possibly dozens of adaptation events (Table S1).

The measles vaccine induces long-lasting immunity ^15,36^. This requires priming of immune cells, T cell responses, and establishment of immune memory ^51^. Our data indicate that the MeV vaccine provides all components for efficient antigen-priming: sensing of viral RNAs leads to upregulation of MHC molecules, especially class I (signal 1), co-stimulatory molecules (signal 2), and T cell-activating cytokines (signal 3).

Our approach utilizing PBMC cultures allowed us to dissect differences in virulence and molecular functions of WT and vaccine MeV and to better understand the molecular basis of MeV vaccine attenuation. In a broader context, this approach is well-suited to address why some live-attenuated vaccines, such as mumps or influenza vaccines, are less efficacious than the measles vaccine and immune responses against these pathogens wane over time ^52^. The key to the success of the measles vaccine could be its high affinity for infecting immune cells and inducing strong innate antiviral responses in them. We may be able to enhance the efficacy and durability of other live-attenuated vaccines by modifying them to replicate the immune-activating features of the measles vaccine.

Limiting to our study are the observed donor-to-donor variations. Especially in our RNA-seq data analysis, we detected strong variations of gene expression levels between the different donors at baseline, likely due to different frequencies of immune cell types in the donor samples. These variations may have masked some gene expression changes induced by the WT, and likely also the C^KO^ virus.

In conclusion, our data provide detailed insights into the molecular mechanisms involved in attenuation of the MeV vaccine. This virus, in contrast to WT strains, elicits strong innate antiviral responses by indiscriminately entering hostile cell types and starting transcription resulting in strong innate antiviral responses. These may be the determinants of the high efficacy of the “self-adjuvanting” MeV vaccine.

## Materials and Methods

### Cell lines and viruses

Vero (ATCC #CCL-81), Vero-hSLAM ^53^, and HEK-293T/17 cells (ATCC #CRL-11268) were cultivated in DMEM supplemented with 10% FBS, 1× L-glutamine, and 1× Penicillin/Streptomycin. Recombinant wild type measles virus strain IC323 expressing GFP [WT: IC323(GFP)_H_] and its C protein-deficient derivative [C^KO^: IC323-C^KO^(GFP)_H_], as well as recombinant vaccine strain measles virus expressing GFP [vac2: vac2(GFP)_H_] were described previously ^33^. Chimeric measles vaccine viruses expressing wild type P/V/C gene [vPwt: vac2(GFP)_H_-P/V/Cwt], wild type H gene [vHwt: vac2(GFP)_h_-Hwt], or both [vPHwt: vac2(GFP)_H_-P/V/Cwt-Hwt] were newly generated in this study. All viruses were rescued by transfecting full-length antigenome-encoding plasmids and helper plasmids into HEK-293T/17 cells as described previously ^54^. Virus stocks were grown in Vero (viruses expressing vaccine strain H) or Vero-hSLAM (viruses expressing wild type strain H), and virus titers were determined by the TCID_50_ method in Vero-hSLAM cells. For all experiments in this study, passage 1 virus stocks were used.

### Cloning of chimeric measles viruses

The original vaccine strain P/V/C was exchanged for wild type P/V/C by three-fragment Gibson Assembly (New England Biolabs # E5510S). For this, WT P/V/C was PCR amplified from p(+)IC323(GFP)_H_ ^7^ with primers PVC_fwd (ATCATTGTTA TAAAAAACTT AGGAACCAGG TCCACACAGC) and PVC_rev (GGAGGCAATC ACTTTGCTCC TAAGTTTTTT ATAATGGATT TAGGTTGTAC TAATTAGGTC GACTGG). The pB(+)MVvac2(GFP)_H_ backbone ^32^ without P/V/C gene was amplified in two fragments with primers vac2_bb1_fwd (CATTATAAAA AACTTAGGAG CAAAGTGATT GCCTC), vac2_bb1_rev (TCCAGTTATC AACGCACTGC TCAT), vac2_bb2_fwd (AGGTGAAGGG TTAACACATG AGCAG), and vac2_bb2_rev (TGTGGACCTG GTTCCTAAGT TTTTTATAAC AATG). The PCR products were purified and used for Gibson Assembly reaction following the manufacturer’s instructions, yielding pB(+)MVvac2(GFP)_H_-P/V/Cwt (vPwt).

The original vaccine strain H gene was exchanged for wild type H gene by restriction/ligation cloning of full length plasmids using PacI/MluI restriction sites, yielding pB(+)MVvac2(GFP)_H_-Hwt (vHwt) and pB(+)MVvac2(GFP)_H_-P/V/Cwt-Hwt (vPHwt).

### PBMC isolation

PBMCs were isolated from apheresis cones from healthy human donors obtained through the German Red Cross Blood Donation Service Hessen-Baden Wuerttemberg (Frankfurt am Main, Germany; samples PEI1-PEI6) or the Mayo Clinic Division of Transfusion Medicine (Rochester, MN, United States; samples MC1-MC13) ^55^. Samples were de-identified (donor age and biological sex were provided by Mayo Clinic, but not German Red Cross) and thus did not require IRB review for research on human subjects according to 45 CFR 46.

Contents of apheresis cones were mixed with PBS and layered over Histopaque-1077 (Sigma-Aldrich, Darmstadt, Germany) or Lymphoprep (Stem Cell Research) density gradient medium according to the manufacturers’ instructions and centrifuged at 800× g for 30 min with acceleration and deceleration set to minimum. The opaque PBMC layer at the interface was collected, mixed with PBS, and pelleted through centrifugation at 500× g, 4°C, for 10 min. Cell pellets were washed once with PBS, then resuspended in 10 ml ACK lysis buffer [150 mM NH_4_Cl; 10 mM KHCO_3_; 0.1 mM EDTA; pH7.2-7.4] and incubated at room temperature for 4 min. The sample volumes were adjusted to 50 ml with PBS, and cells were pelleted again by centrifugation. Pellets were washed once with PBS and then resuspended in 10 ml RPMI supplemented with 10% FBS, 1× L-glutamine, and 1× Penicillin/Streptomycin (RPMI+). Cells were counted at 1:100 dilutions with trypan blue using an Improved Neubauer hemocytometer.

Cell concentrations were then adjusted to 1-2×10^7^ cells/ml and samples were supplemented with PHA-M at a final concentration of 10 μg/ml. Cell suspension were incubated overnight in upright standing tissue culture flasks at 37°C and 5% CO_2_.

### PBMC infections

Infections were carried out in PBMCs after overnight culture in PHA-M-containing RPMI+, which leads to upregulation of the MeV entry receptor CD150/SLAM in T cells. An adequate volume of cell suspension was removed from the culture flasks, cells were pelleted by centrifugation (500× g, 4°C, 10 min), pellets were resuspended in 5-10 ml PBS, and cells were counted using an Improved Neubauer hemocytometer. 1-5×10^6^ cells were transferred into 15 ml tubes and pelleted again by centrifugation. Pellets were resuspended in OptiMEM containing virus inoculum for the targeted multiplicity of infection of 0.1, and the total volume was adjusted with OptiMEM to obtain a cell density of 4×10^6^ cells/ml. Uninfected controls were incubated in OptiMEM at the same density. Cells were incubated at 37°C for 2 h and resuspended by shaking every 20 min. Afterwards, cells were pelleted by centrifugation (500x g, 4°C, 10 min), washed once with PBS, resuspended in RPMI+ with 10 μg/ml PHA-M at a cell density of 1.0×10^7^ cells/ml, seeded in round-bottom 96-or flat-bottom 24-wells depending on the total sample volume and incubated at 37°C and 5% CO_2_.

Table S3 summarizes the different viruses studied and assays performed with the PBMCs from individual donors.

### Virus growth kinetics in PBMCs

To determine infectious virus titers at different timepoints post infection, PBMCs were carefully resuspended in their culture supernatant, and a volume equivalent of 2.5×10^5^ cells was removed and diluted 10-fold in DMEM without supplements. From this dilution, a 10-fold dilution series was created (total of 8 dilutions). 50-100 µl of each dilution was added to 96-wells containing 100-150 µl of Vero-hSLAM cell suspensions in DMEM supplemented with 10% FBS, 1× L-glutamine, and 1× Penicillin/Streptomycin. Each dilution was added to a total of 4 replicates. Vero-hSLAM cells were incubated at 37°C and 5% CO_2_ for 4 days, after which viral cytopathic effect (CPE) in form of destroyed cell layer or syncytia formation was determined for each well. Virus titers were determined through equation 1:

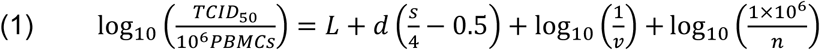

with:

*L*: last dilution in which all four replicates were CPE-positive

*d*: −log_10_ of dilution factor (here: *d* = 1)

s: number of CPE-positive wells starting at the last dilution in which all four replicates were CPE-positive

*v*: inoculation volume in ml

*n*: number of cells removed from culture (here: *n* = 2.5×10^5^)

### Flow cytometry analysis

1.0-2.5×10^6^ infected or uninfected PBMCs were collected in flow cytometry tubes 48 h post infection by passing through a 35 μm cell strainer. Cells were washed twice with PBS and pelleted through centrifugation at 500 × g, 4°C, for 10 min. Cells were stained with Zombie NIR viability dye (1:1000; Biolegend #423106) in PBS at room temperature in the dark for 15 min, followed by two wash steps with PBS. Surface markers were stained using a cocktail of antibodies listed below. Antibodies were diluted to the indicated final dilutions in PBS supplemented with Human TruStain FcX (1:100; Biolegend #422302) and True-Stain Monocyte Block (1:100; Biolegend #426102). Cell pellets were resuspended in 100 μl antibody cocktail and incubated at 4°C in the dark for 45 min. Cells were washed twice with PBS, and then fixed with 250 μl Cytofix Fixation Buffer (BD #554655) at 4°C in the dark for 15 min. After two more washes with PBS, cells were resuspended in 250 μl Cell Staining Buffer (Biolegend #420201) supplemented with Tandem Stabilizer (1:1000; Biolegend #421802). Samples were analyzed on a 4-color (V/B/YG/R) Cytek Aurora Spectral Flow Cytometer within 24 hours of staining.

Unstained and single-color controls on PBMCs and Ultracomp eBeads (Thermo Fisher #01-2222-42) for unmixing were prepared accordingly and stored in the reference library. Unmixed data was analyzed using FlowJo Portal (BD, v.10.9.0 CL) and visualized with Prism (GraphPad, v.10.4.2).

### Western blot analysis

1×10^7^ infected or uninfected PBMCs were collected 48 h post infection and pelleted by centrifugation at 500× g, 4°C, for 10 min. Cells were washed once with PBS. Cell pellets were resuspended in 100 μl protein lysis buffer [10 mM HEPES, pH 7.9; 200 mM NaCl; 5 mM KCl; 10% (v/v) glycerol; 0.5 % (v/v) NP-40; 1 mM EDTA; 1 mM Na-orthovanadate; 5 mM NaF; 1 mM PMSF; 1 mM DTT; 1% (v/v) Protease inhibitor cocktail (Sigma Aldrich #P8340); 1% (v/v) Phosphatase inhibitor cocktail 3 (Sigma Aldrich #P0044)] and incubated on ice for 15 min.

Nuclei and cell debris were pelleted by centrifugation at 21000× g, 4°C, for 30 min, and cleared supernatants were collected and stored at −20°C. For SDS-PAGE, equal volumes of lysates and 2x urea sample buffer [200 mM Tris, pH 6.8; 8 M urea; 5 % (w/v) SDS; 0.1 mM EDTA; 0.03% (w/v) bromophenol blue; 1.5% (w/v) DTT] were mixed and heated at 95°C for 5-10 min. 10 μl of lysates were loaded in each lane of 3.9%/10% polyacrylamide gels. Samples were separated at 100-120 V for 1-1.5 h. Proteins were transferred to Immobilon-P PVDF membranes (Sigma Aldrich #IPVH00010) at 400 mA and 4°C using wet transfer. Membranes were blocked with 5% (w/v) BSA in 1×TBS for 1 h at room temperature. Membranes were incubated with primary antibody solutions overnight at 4°C, then washed three times with 1× TBS +0.05% Tween-20 (1×TBST). Membranes were incubated with HRP-conjugated secondary antibodies in 1×TBST for 1 h at room temperature, then washed three times with 1×TBST. For β-actin loading control, membranes were incubated with HRP-conjugated anti-hu-β-actin antibody diluted in 1×TBST for 30 min at room temperature. For development, membranes were soaked with SuperSignal West Pico PLUS Chemiluminescent Substrate (Thermo Fisher #34580) for 5-10 min and images at multiple exposure times were obtained with a ChemiDoc Imager (Biorad). Images were quantified using Image Lab (v.6.0, Biorad) and cropped with Adobe Photoshop (v.24.6.0).

### Antibodies

The following antibodies were used for flow cytometry staining:

BV421 anti-hu-CD123 (1:100; clone 6H6; Biolegend #306018); PacificBlue anti-hu-CD20 (1:100; clone 2H7; Biolegend #302320); BV480 anti-hu-CD38 (1:100; clone HIT2; BD #566137); BV510 anti-hu-CD27 (1:100; clone O323; Biolegend #302836); BV605 anti-hu-CD11b (1:50; clone ICRF44; Biolegend #301332); BV650 anti-hu-CD45RA (1:100; clone HI100; Biolegend #304136); BV711 anti-hu-CD56 (1:50; clone 5.1H11; Biolegend #362542); BV750 anti-hu-CD16 (1:100; clone 3G8; Biolegend #302082); BV785 anti-hu-CD127 (1:50; clone A019D5; Biolegend #351330); SparkBlue550 anti-hu-IgM (1:50; clone MHM-88; Biolegend #314555); SparkBlue574 anti-hu-CD4 (1:50; clone SK3; Biolegend # 344680); PerCP anti-hu-CD45 (1:50; clone 2D1; Biolegend #368506); BB700 anti-hu-CD25 (1:100; clone M-A251; BD #566447); RB780 anti-hu-CD8 (1:400; clone RPA-T8; BD #568684); PE anti-hu-CD150 (1:50; clone A12(7D4); Biolegend #306308); PE-Dazzle594 anti-hu-CD197 (1:50; clone G043H7; Biolegend #353236); PE-Cy5 anti-hu-CD11c (1:50; clone 3.9; Biolegend #301610); PE-Fire700 anti-hu-TCRγδ (1:50; clone B1; Biolegend #331238); APC anti-hu-HLA-DR (1:100; clone L243; Biolegend #307610); SparkNIR685 anti-hu-CD3 (1:200; clone SK7; Biolegend #344862); AlexaFluor700 anti-hu-IgD (1:200; clone W18340F; Biolegend #307820); APC-Fire810 anti-hu-CD14 (1:50; clone 63D3; Biolegend #367156).

The following primary antibodies were used for western blot analyses:

rabbit anti-hu-IRF3 (1:1000; clone D6I4C; Cell Signaling #11904); rabbit anti-hu-phospho-IRF3_S396_ (1:1000; clone 4D4G; Cell Signaling #4947); rabbit anti-hu-IRF7 (1:1000; clone D2A1J; Cell Signaling #13014); rabbit anti-hu-phospho-IRF7_S477_ (1:1000; clone D7E1W; Cell Signaling #12390); rabbit anti-hu-STAT1 (1:1000; clone D1K9Y; Cell Signaling #14994); rabbit anti-hu-phospho-STAT1_Y701_ (1:1000; clone D4A7; Cell Signaling #7649); rabbit anti-hu-STAT2 (1:1000; clone D9J7L; Cell Signaling #72604); rabbit anti-hu-phospho-STAT2_Y690_ (1:1000; clone D3P2P; Cell Signaling #88410); rabbit anti-hu-PKR (1:1000; clone D7F7; Cell Signaling #12297); rabbit anti-hu-phospho-PKR_T446_ (1:1000; clone E120; Abcam #ab32036); rabbit anti-MeV-N_505_ (1:5000) ^33,34,56^; HRP-conjugated mouse anti-hu-β-actin (1:20000; clone AC-15; Sigma Aldrich #A3854).

The following secondary antibody was used for western blot analyses:

HRP-conjugated AffiniPure goat anti-rabbit IgG H+L (1:20000; Jackon ImmunoResearch #111-035-144).

### IFNβ enzyme-linked immunosorbent assay

Supernatants from uninfected or infected PBMCs were harvested at 48 h post infection by pelleting cells at 500× g, 4°C, for 10 min. Supernatants were carefully removed and stored at −80°C until assay was performed. A commercial Human IFNβ ELISA Kit (Bio-Techne #DIFNB0) was used to quantify levels of IFNβ in supernatants strictly following the instructions in the manual.

### RNA extraction

4×10^6^ cells per condition and donor were collected at 48 h post infection and washed once with PBS. Cells were pelleted by centrifugation at 500× g, 4°C, for 10 min. Total RNA was isolated using TRIzol (Thermo Fisher #15596026) according to the manufacturer’s instructions. The resulting RNA pellet was reconstituted in 30 μl DEPC-treated H_2_O, and samples were stored at −80°C.

### RNA-seq library preparation and sequencing

Stranded total RNA-seq libraries were generated and sequencing was performed by Azenta/Genewiz (Leipzig, Germany). Ribosomal and globin RNAs were depleted prior to library preparation. Each sample was sequenced to a depth of at least 5×10^7^ reads on an Illumina NextSeq platform using 2× 150 bp paired-end sequencing following a protocol for stranded total RNA libraries.

### RNA-seq data alignment

All bioinformatic operations were conducted on the Galaxy Project platform (v.23.1.rc1)^57^. Raw reads were preprocessed with fastp (v.0.23.2) ^58^ to filter low quality reads and trim adapter sequences using default settings. Quality assessment was performed using MultiQC (v.1.11) ^59^ using the JSON outputs of fastp. Reads were mapped to the human reference genome (GRCh38.p14 (GCA_000001405.15) with the splice-aware STAR mapping algorithm (v.2.7.10b) ^60^ using default settings for strand-specific, paired-end library sequencing. Mapped reads were quantified with the built-in gene quantification (GeneCounts) using the GTF annotation of GRCh38 (v.42, 07/20/2022).

### Differential gene expression analysis

Differential gene expression in virus-infected samples over mock infection was assessed using DESeq2 (v.1.34.0) ^61^ using the raw counts from STAR GeneCounts as input. A pre-filter for minimal gene counts was set to 5 and data shrinkage was activated. The four different infection conditions (UI, WT, C^KO^, and vac2) were treated as primary factors, and donors were treated as secondary factors to reduce donor-specific effects. Principal component plots (PC-plots) were generated with DESeq2. Significantly deregulated genes were those with −log_10_(adjusted P-value) ≥1.301029 and log_2_(fold-change) ≥±0.584962. To generate a z-score plot, the DESeq2 normalized count data of all conditions for genes that were significantly deregulated in infection with the WT, C^KO^ and vac2 were combined and the heatmap2 tool (v.3.1.3) was used for plotting. Data transformation was disabled, the z-score was calculated on rows (normalized gene counts) and clustering was performed on rows and columns.

### Viral mRNA gradient analysis

Viral reads were aligned to the (+)strand of the MeV reference genomes using HISAT2 (v.2.2.1) ^62^ with skip reverse strand of reference activated, and read counts per MeV gene were quantified using the GeneCounts feature of STAR (v.2.7.10b) ^60^ using GTF files defining the MeV gene regions. Read counts were used to calculate transcripts per million (TPM) values to obtain expression values corrected for gene length and total sequencing depth following equation 2:

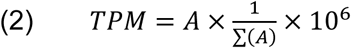

with *A* = (raw counts per gene) / (gene length in kilobases).

### Gene ontology analysis

Functional enrichment of significantly deregulated genes was performed using ShinyGO (v.0.82) ^63^. Gene sets of up- and down-regulated genes were tested separately for the gene ontology (GO) term biological process (BP). Type-I IFN-stimulated genes (ISGs) were identified in the entire gene lists returned from DESeq2 using the Interferome database (v.2.0) ^35^.

### Statistical analysis

In all graphs, data derived from individual donors are specified by unique symbols as provided in Supplemental Table 1 (exception: RNA-seq data). Graphs show individual values, mean values, and standard deviation if not stated differently in the figure legend. Statistical tests were performed with Prism (GraphPad, v.10.4.1). Viral growth kinetics were analyzed using repeated measures two-way ANOVA with Geisser-Greenhouse correction followed by Tukey’s multiple comparisons test with individual variances computed for each comparison. Flow cytometry and IFNβ ELISA data were analyzed by repeated measure one-way ANOVA, or mixed-effects analysis, with Geisser-Greenhouse correction followed by Tukey’s multiple comparisons test with individual variances computed for each comparison. Statistical analysis of RNA-seq data was performed by DESeq2 ^61^ and ShinyGO^63^.

## Acknowledgments

The authors dedicate this study to Karl-Klaus Conzelmann (1955-2025), who pioneered the field of negative strand RNA virus reverse genetics and innate immune evasion. We would like to thank the Mayo Clinic Blood Component Laboratory for excellent support with donor material.

## Author contributions

Conceptualization: FGMA, OS, KA, RC, BS, CKP

Methodology: FGMA, SP, KA, BS, CKP

Investigation: FGMA, SP, FMA, DAS, OS, CKP

Visualization: FGMA, SP, FMA, OS, CKP

Funding acquisition: CKP, BS

Project administration: CKP, BS

Supervision: CKP, BS, OS

Writing – original draft: FGMA, OS, KA, RC, BS, CKP

Writing – review & editing: FGMA, SP, FMA, DAS, OS, KA, RC, BS, CKP

## Diversity, equity, ethics, and inclusion

PBMCs from human donors were obtained through the German Red Cross Blood Donation Service Baden-Württemberg-Hessen (Frankfurt am Main, Germany) and through the Mayo Clinic Division of Transfusion Medicine (Rochester, MN, United States). Studies were performed with anonymized samples and thus were exempt from IBC regulations. Female and male donors were equally represented.

## Data and materials availability

Raw and analyzed RNAseq data will be made available in the Gene Expression Omnibus (GEO) database upon publication in a peer-reviewed journal. Other data are available in the main text or the supplementary materials. Viruses and other materials generated in this study will be made available under institutional material transfer agreements upon request to the corresponding author.

## Supplementary Figures and Tables

**Fig. S1.**
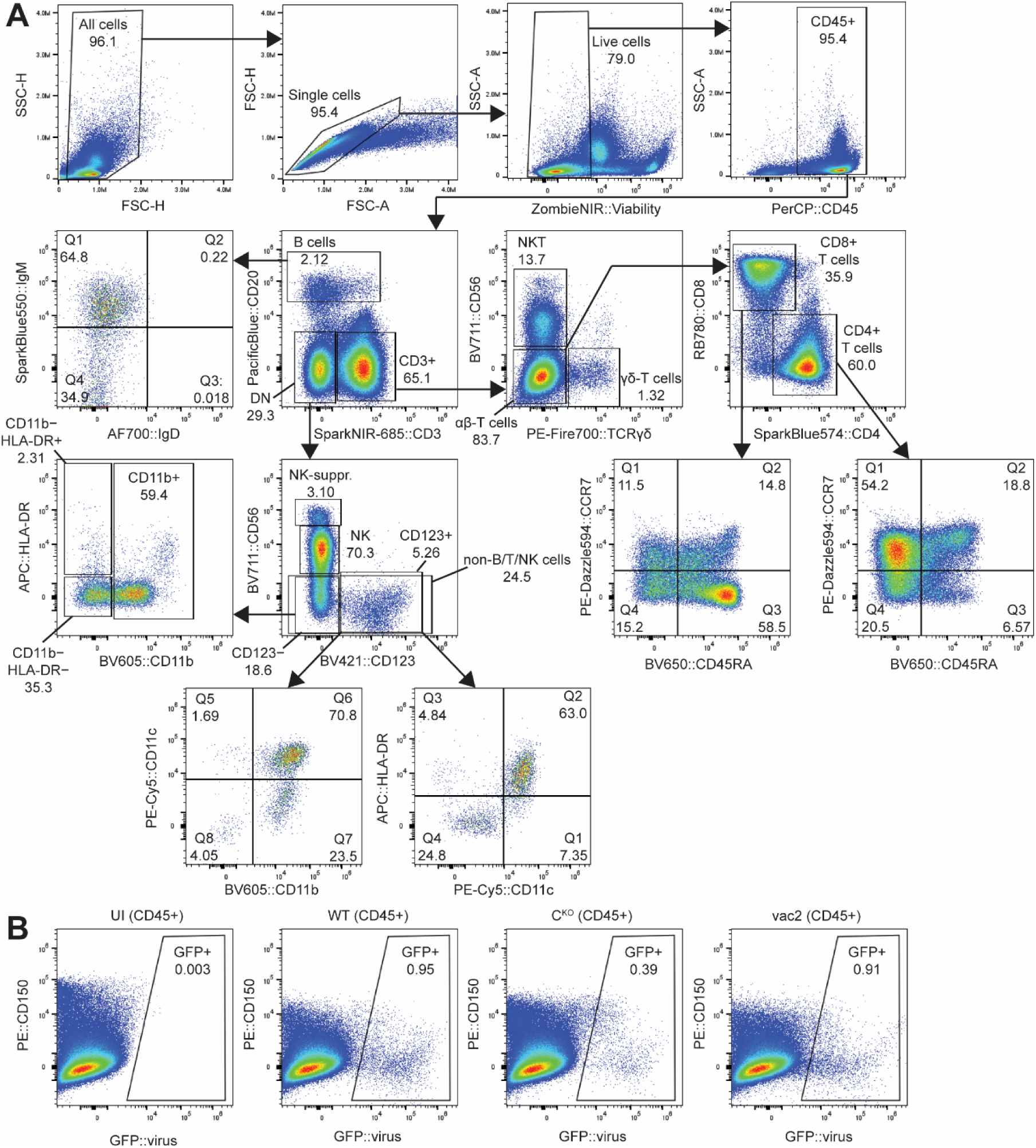
Spectral flow cytometry gating strategy for human PBMCs. (**A**) Flow chart of gating to analyze different immune cell types in PBMCs. (**B**) Representative pseudocolor plots of CD150 expression against GFP expression (infection marker) in total live CD45+ cells from one donor.

**Fig. S2.**
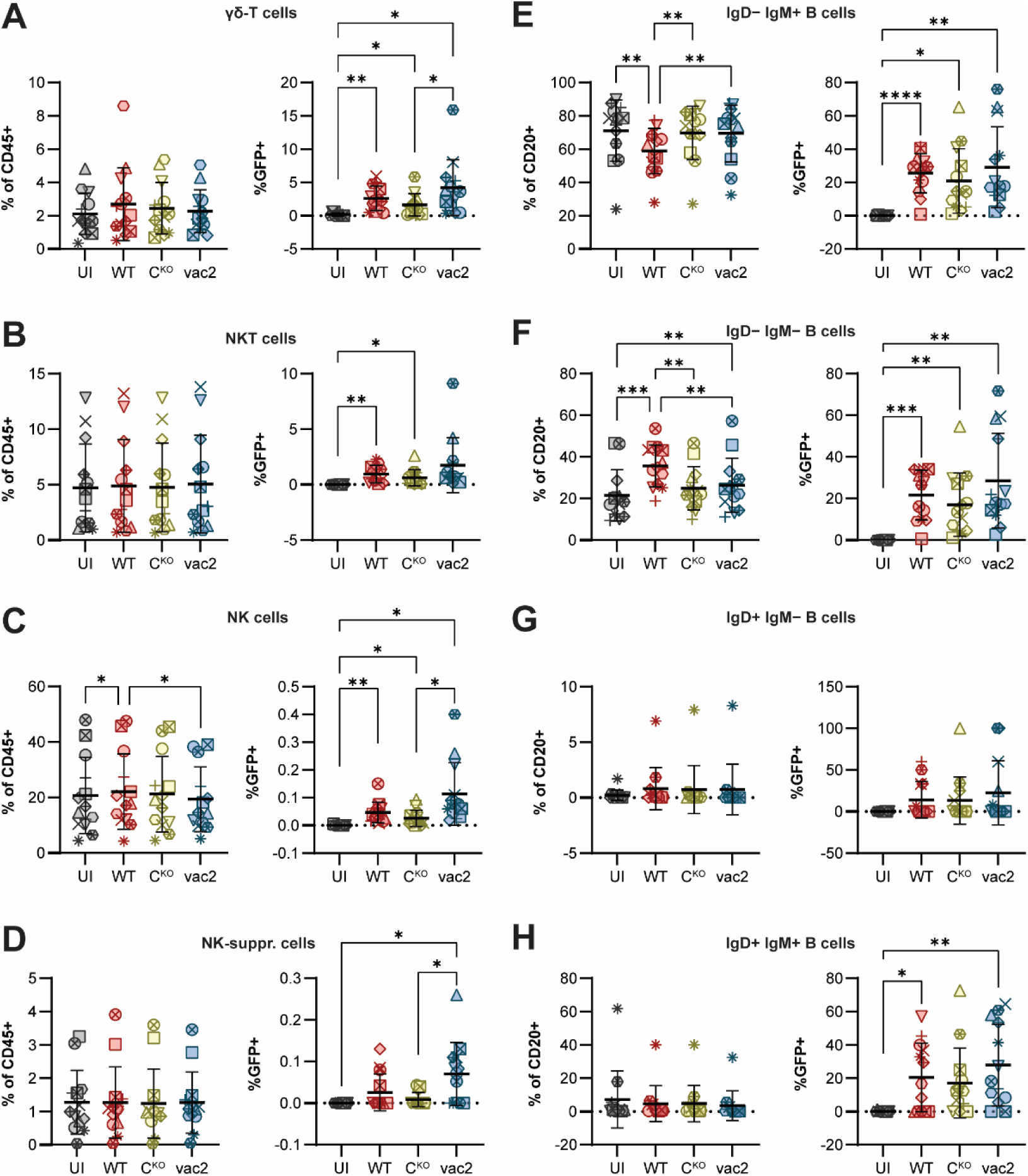
Flow cytometry quantification of T cell subsets. Flow cytometry analysis of infection rates based on GFP-reporter expression in different immune cell subsets from human donors (n = 13). Statistics: Repeated measures one-way ANOVA with Tukey’s multiple comparisons test. (**A**)-(**D**) Left diagrams show cell type frequencies in CD4+ population; right diagrams show GFP+ cells of CD4+ T cell subtype. (A) Naïve CD4+ T cells. (**B**) Central memory (CM) CD4+ T cells. (**C**) Effector memory (EM) CD4+ T cells. (**D**) Terminal effector memory (TEMRA) CD4+ T cells. (**E**)-(**H**) Left diagrams show cell type frequencies in CD8+ population; right diagrams show GFP+ cells of CD8+ T cell subtype. (**E**) Naïve CD8+ T cells. (**F**) Central memory (CM) CD8+ T cells. (**G**) Effector memory (EM) CD8+ T cells. (**H**) Terminal effector memory (TEMRA) CD8+ T cells. In all graphs, each donor is identified with a unique symbol. Bars indicate mean values and error bars show standard deviations. Asterisks indicate statistical significance levels: * P≤0.05; ** P≤0.01; *** P≤0.001.

**Fig. S3.**
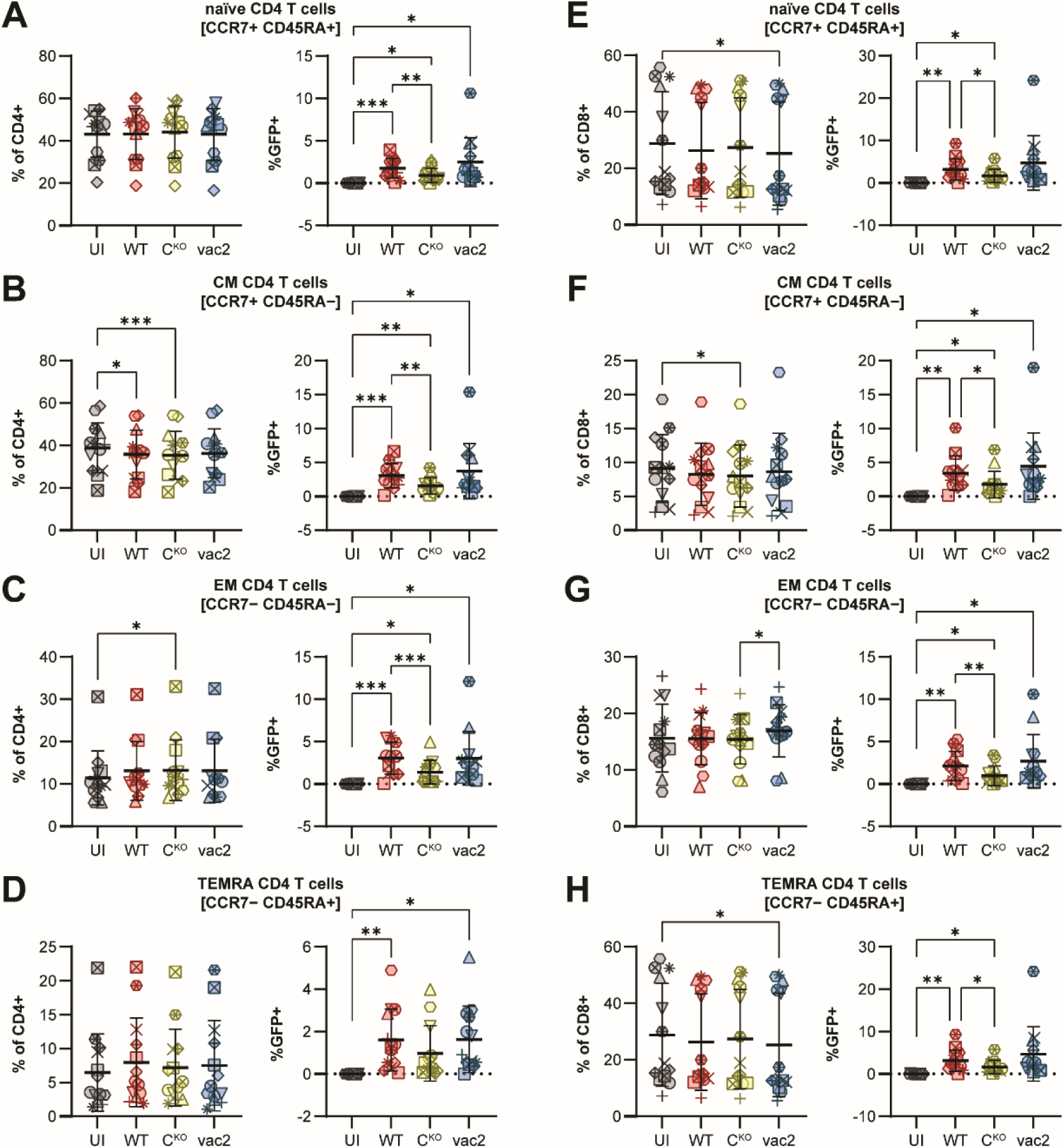
Flow cytometry quantification of additional immune cell types and B cell subsets. Flow cytometry analysis of infection rates based on GFP-reporter expression in different immune cell subsets from human donors (n = 13). Statistics: Repeated measures one-way ANOVA with Tukey’s multiple comparisons test. (**A**)-(**D**) Left diagrams show cell type frequencies in CD45+ population; right diagrams show GFP+ cells of cell type. (**A**) γδ-T cells. (**B**) NKT cells. (**C**) NK cells. (**D**) NK suppressor cells. (**E**)-(**H**) Left diagrams show cell subtype frequencies in CD20+ (B cell) population; right diagrams show GFP+ cells of B cell subtype. (**E**) IgD− IgM+ B cells. (**F**) IgD− IgM− B cells. (**G**) IgD+ IgM− B cells. (**H**) IgD+ IgM+ B cells. In all graphs, each donor is identified with a unique symbol. Bars indicate mean values and error bars show standard deviations. Asterisks indicate statistical significance levels: * *P*≤0.05; ** *P*≤0.01; *** *P*≤0.001.

**Fig. S4.**
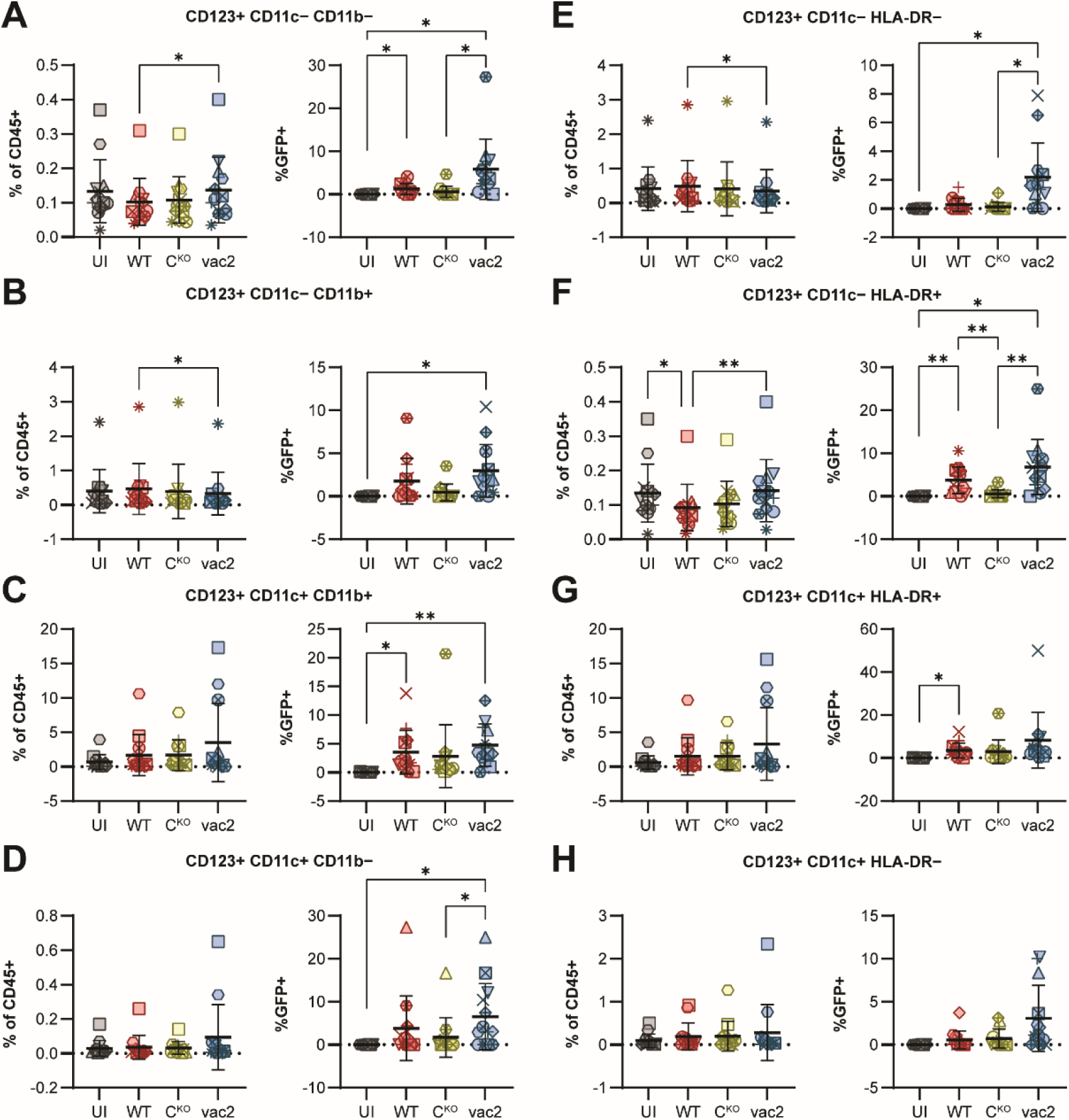
Flow cytometry quantification of CD123+ non-B/T/NK cell subsets. Flow cytometry analysis of infection rates based on GFP-reporter expression in different immune cell subsets from human donors (n = 13). Statistics: Repeated measures one-way ANOVA with Tukey’s multiple comparisons test. (**A**)-(**H**) Left diagrams show cell type frequencies in CD45+ population; right diagrams show GFP+ cells of CD123+ non-B/T/NK cell subtype. (**A**) CD123+ CD11c− CD11b− cells. (**B**) CD123+ CD11c− CD11b+ cells. (**C**) CD123+ CD11c+ CD11b+ cells. (**D**) CD123+ CD11c+ CD11b− cells. (**E**) CD123+ CD11c− HLA-DR− cells. (**F**) CD123+ CD11c− HLA-DR+ cells. (**G**) CD123+ CD11c+ HLA-DR+ cells. (**H**) CD123+ CD11c+ HLA-DR− cells. In all graphs, each donor is identified with a unique symbol. Bars indicate mean values and error bars show standard deviations. Asterisks indicate statistical significance levels: * P≤0.05; ** P≤0.01.

**Fig. S5.**
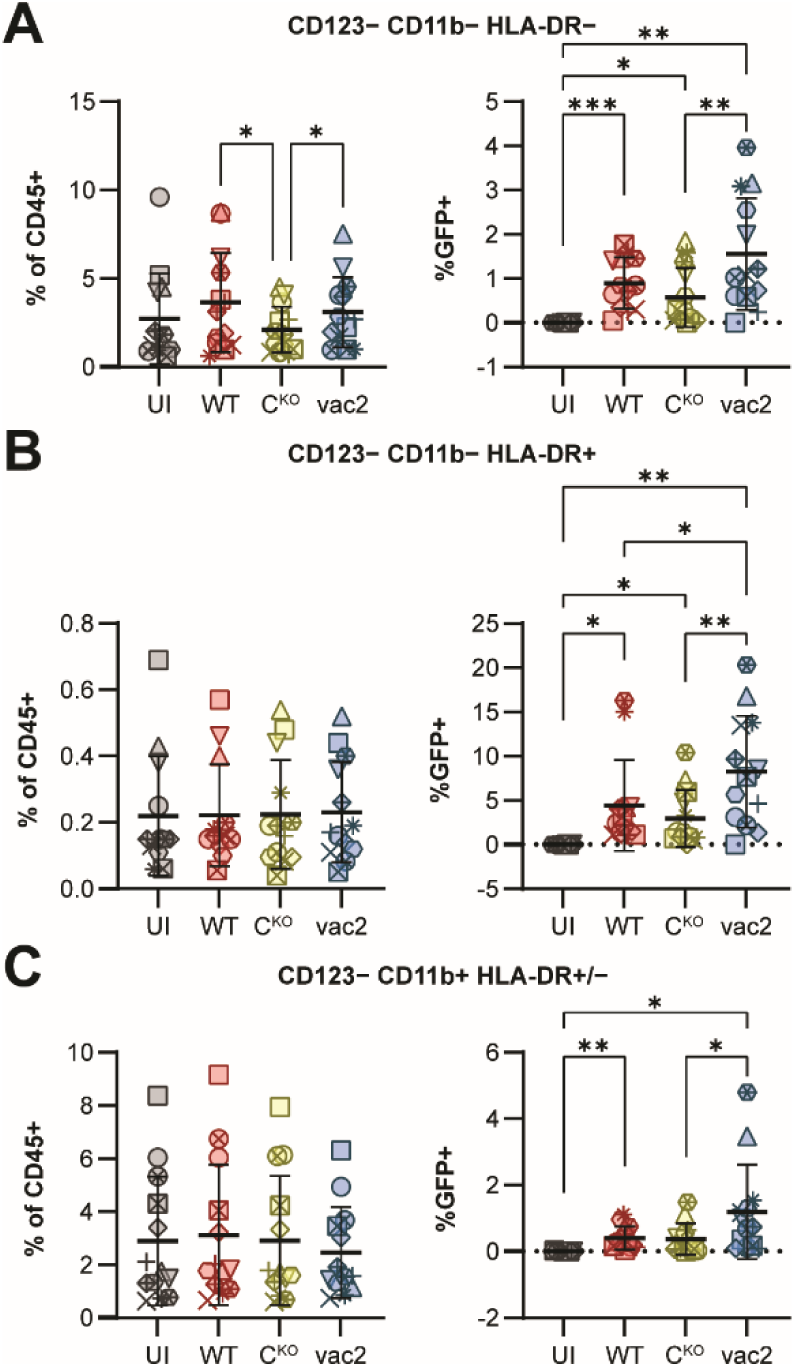
Flow cytometry quantification of CD123− non-B/T/NK cell subsets. Flow cytometry analysis of infection rates based on GFP-reporter expression in different immune cell subsets from human donors (n = 13). Statistics: Repeated measures one-way ANOVA with Tukey’s multiple comparisons test. (**A**)-(**C**) Left diagrams show cell type frequencies in CD45+ population; right diagrams show GFP+ cells of CD123− non/B/T/NK cell subtype. (**A**) CD123− CD11b− HLA-DR− cells. (**B**) CD123− CD11b− HLA-DR+ cells. (**C**) CD123− CD11b+ HLA-DR+/− cells. In all graphs, each donor is identified with a unique symbol. Bars indicate mean values and error bars show standard deviations. Asterisks indicate statistical significance levels: * P≤0.05; ** P≤0.01; *** P≤0.001.

**Fig. S6.**
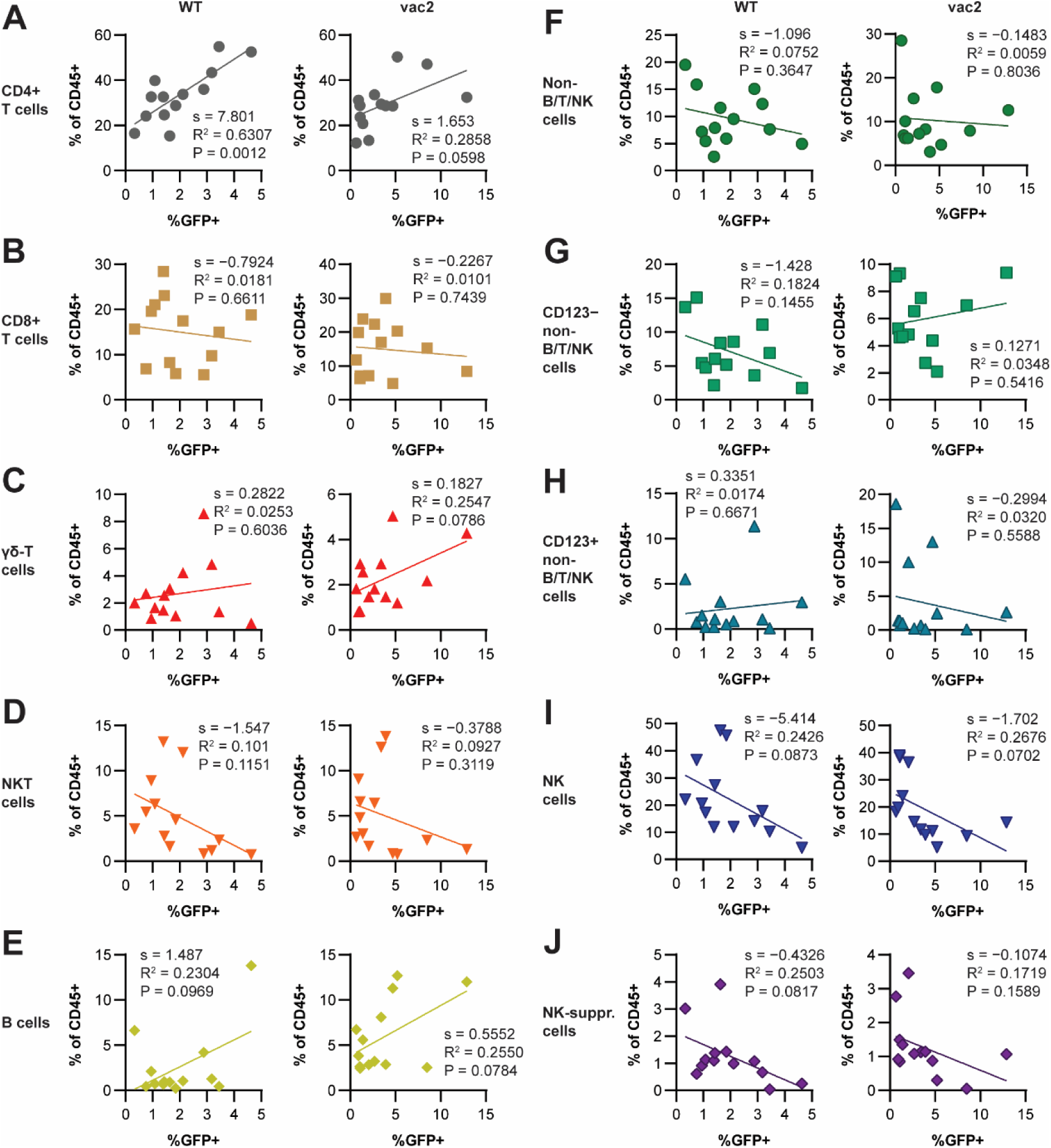
Correlation analysis of immune cell type frequencies and MeV infectivity. Simple linear regression analysis of virus (WT: left diagrams; vac2: right diagrams) infection rate (%GFP+) in CD45+ cells with cell type frequencies in CD45+ population. Each symbol represents one donor (n = 13). Values calculated with GraphPad Prism: s (slope of fitted line), R^2^ (fit), P (p value). (**A**) CD4+ T cells. (**B**) CD8+ T cells. (**C**) γδ-T cells. (**D**) NKT cells. (**E**) B cells. (**F**) Non-B/T/NK (CD3− CD20− CD56−) cells. (**G**) CD123− non-B/T/NK cells. (**H**) CD123+ non-B/T/NK cells. (**I**) NK cells. (**J**) NK suppressor cells.

**Fig. S7.**
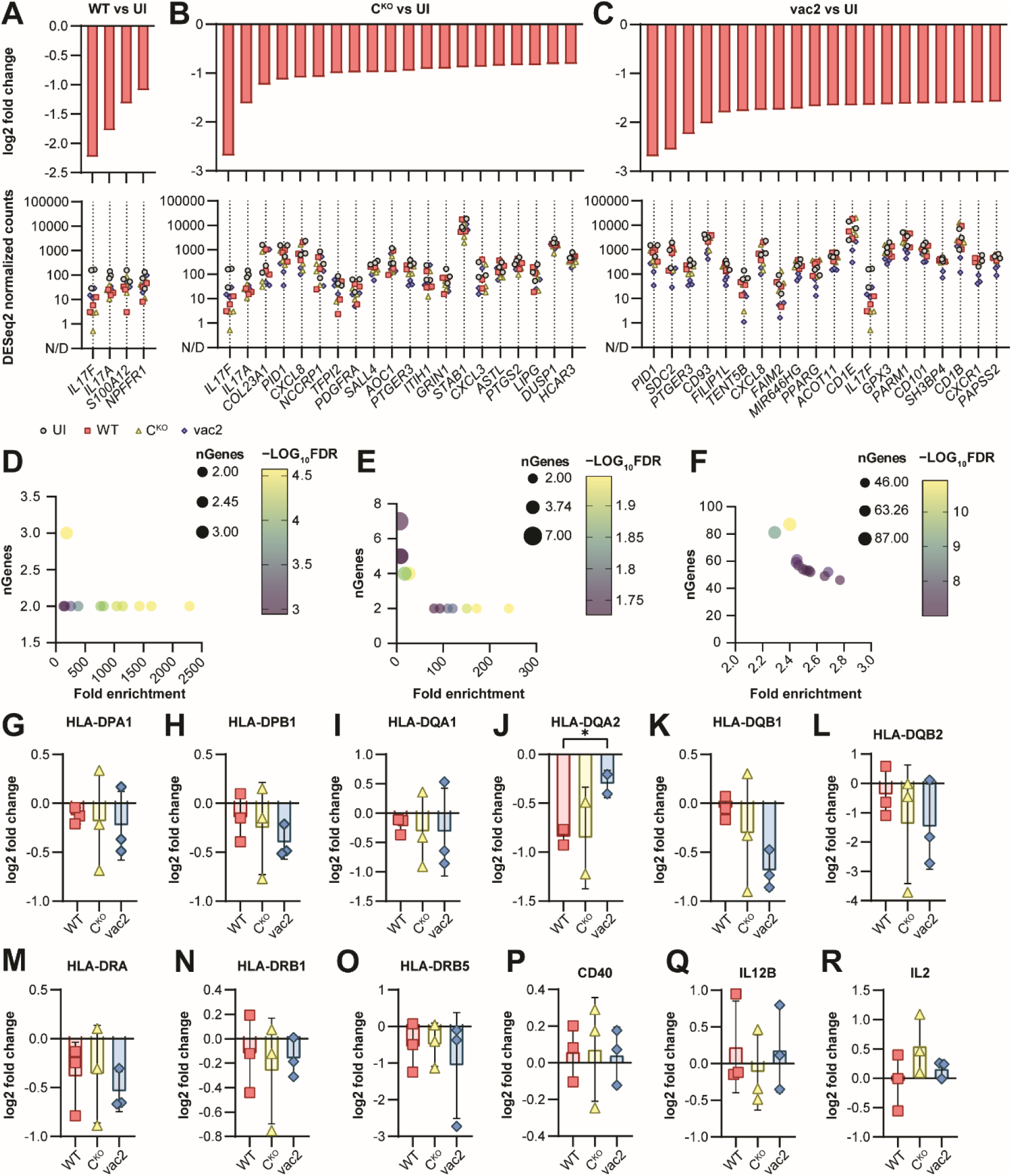
Analysis of downregulated genes after MeV infection. (**A**)-(**C**) Top downregulated genes during infection, ranked in ascending order of log2(fold change) (top diagrams). The bottom diagrams indicate the normalized read counts of each individual sample. (**A**) WT infection. (**B**) C^KO^ infection. (**C**) vac2 infection. (**D**)-(**F**) GO term analysis of genes upregulated during infection. Circle color indicates −log10(FDR) determined by ShinyGO. Circle size is proportional to the number of genes (nGenes) identified by ShinyGO. (**D**) WT infection. (**E**) C^KO^ infection. (**F**) vac2 infection. (**G**)-(**R**) Log2 fold change of mRNA levels of virus-infected samples compared to uninfected derived from RNA-seq data for genes involved in T cell-responses. (**G**)-(**O**) MHC-II. (**P**) Co-stimulatory molecule CD40. (**Q**)-(**R**) Signal 3 cytokines. Statistics: repeated measures one-way ANOVA with Tuckey’s multiple comparisons test. Bars indicate mean values and error bars show standard deviations. Asterisks indicate statistical significance levels: * *P*≤0.05.

**Fig. S8.**
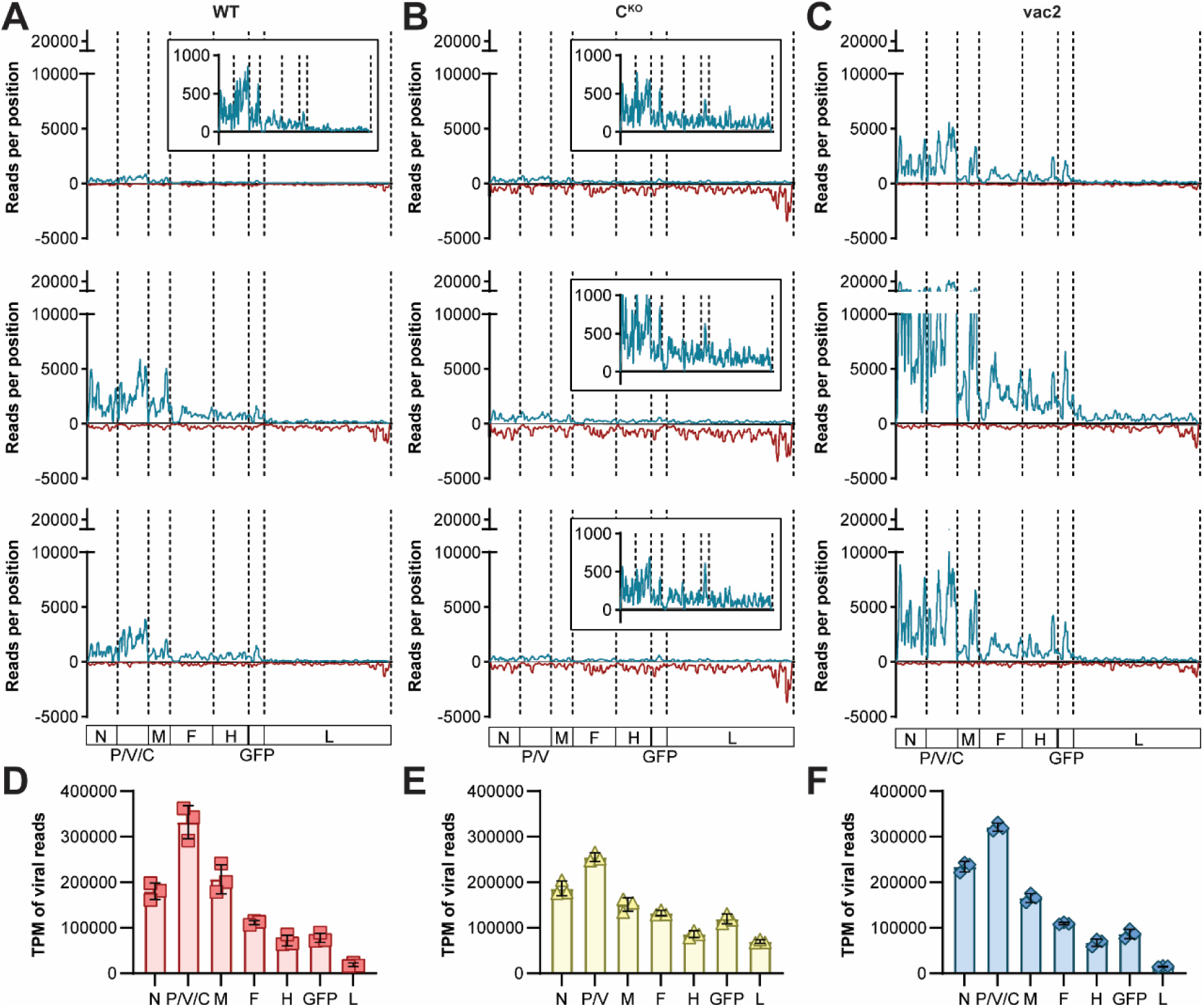
Viral genome coverage in RNA-seq datasets. (**A**)-(**C**) Coverage plots of reads mapped to the MeV reference genomes (x-axis). Coverage of (+)strand reads (=mRNA, antigenome) is depicted as positive value (blue lines) and coverage of (−)strand reads (=genome) is depicted as negative value (red lines). Inserts are zoomed-in views in case of low (+)strand coverage. Vertical dotted lines indicate gene borders. (**A**) WT infected samples. (**B**) C^KO^ infected samples. (**C**) vac2 infected samples. (**D**)-(**F**) Viral mRNA transcription gradients based on calculated transcripts per million (TPM) values for each viral gene. Only (+)strand-aligned reads were included. (**D**) WT infection. (**E**) C^KO^ infection. (**F**) vac2 infection.

**Fig. S9.**
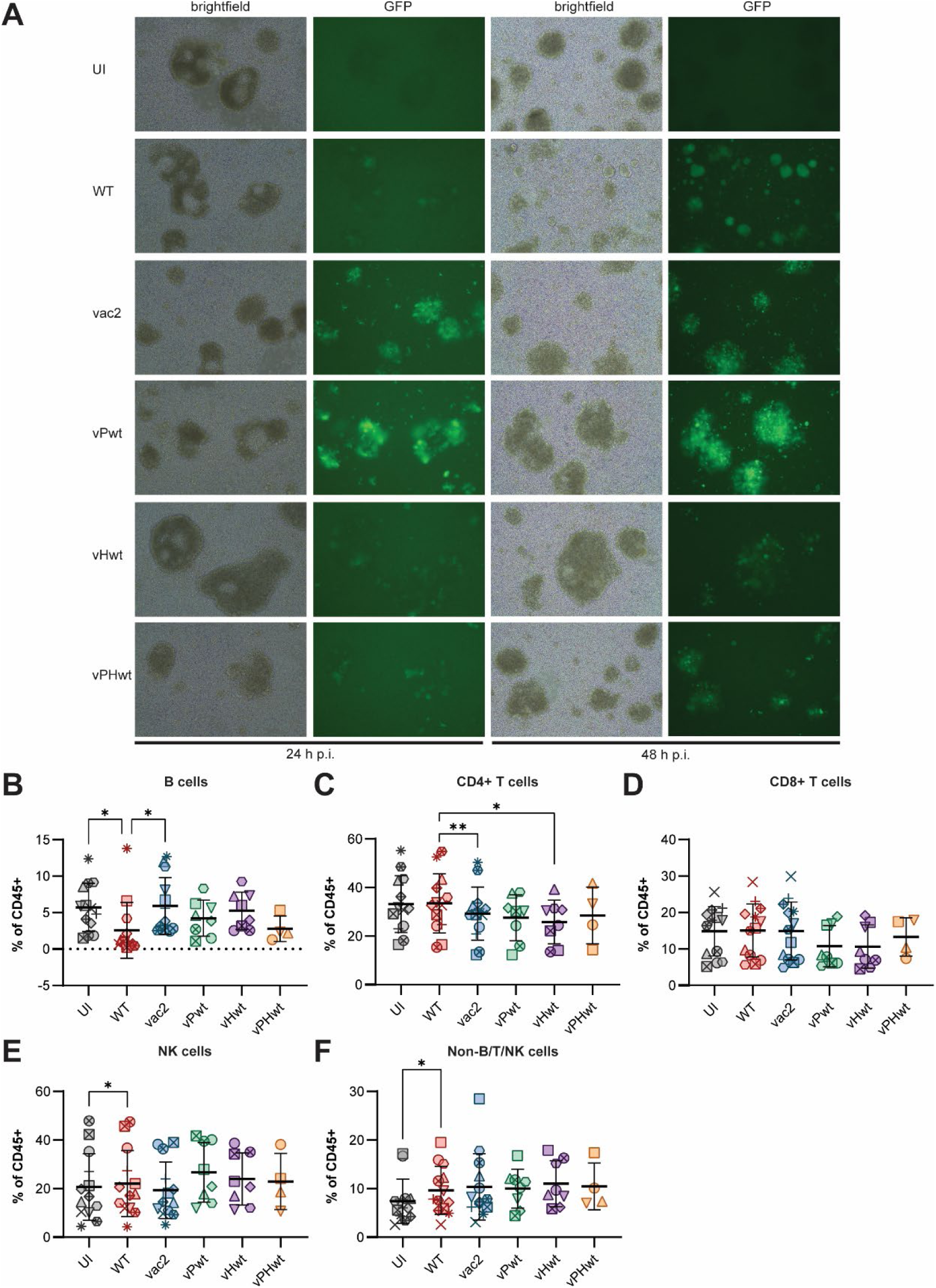
Infection of PBMCs with chimeric MeVs. (**A**) Brightfield images and GFP fluorescence of PBMCs infected with indicated viruses or left uninfected (UI) 24 and 48 h post infection. Images were taken with 10X objective. (**B**)-(**F**) Cell type frequencies of different immune cell subsets from human donors (WT, vac2: n = 13; vPwt, vHwt: n = 8; vPHwt: n = 4). Statistics: Repeated measures one-way ANOVA with Tukey’s multiple comparisons test. Asterisks indicate statistical significance levels: * P≤0.05; ** P≤0.01.

**Fig. S10.**
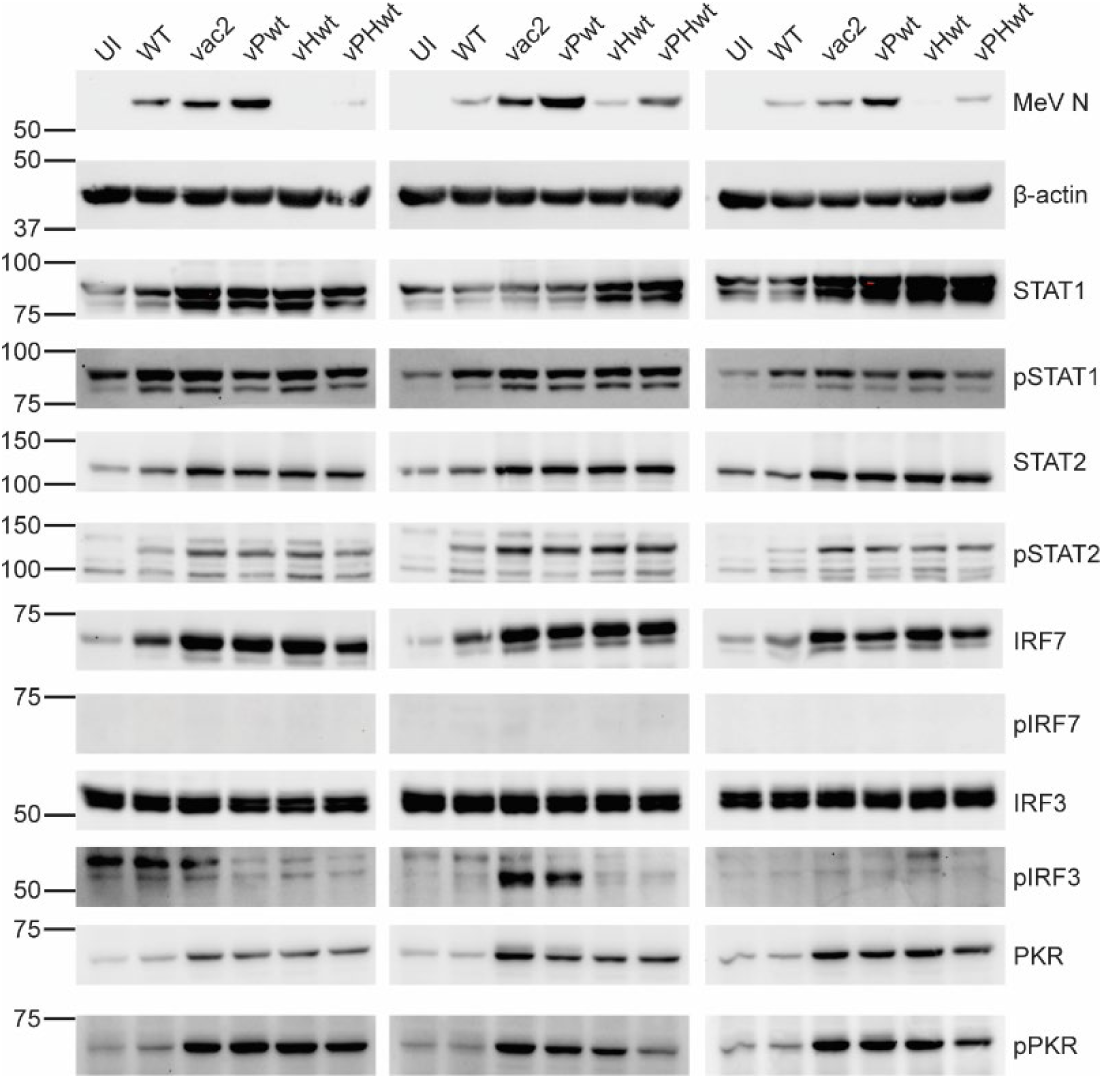
Western blot analysis of MeV infected PBMCs. Three additional donors included in quantifications in Fig. 5C-J.

**Table S1.**
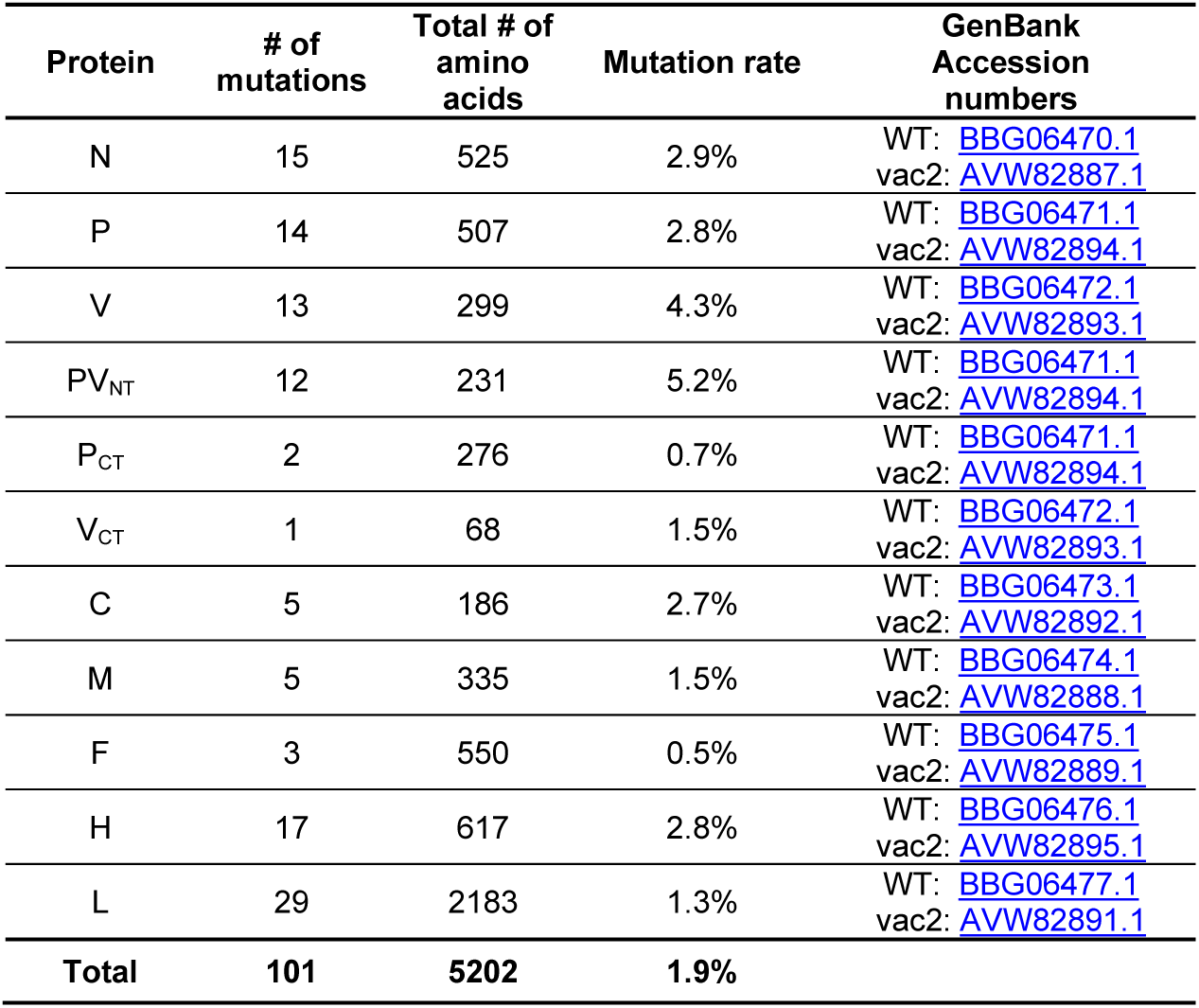
Number of amino acid differences between WT (IC323) and vaccine (vac2) MeV proteins.

| Protein | # of mutations | Total # of amino acids | Mutation rate | GenBank Accession numbers |
| --- | --- | --- | --- | --- |
| N | 15 | 525 | 2.9% | WT: BBG06470.1<br>vac2: AVW82887.1 |
| P | 14 | 507 | 2.8% | WT: BBG06471.1<br>vac2: AVW82894.1 |
| V | 13 | 299 | 4.3% | WT: BBG06472.1<br>vac2: AVW82893.1 |
| PV<sub>NT</sub> | 12 | 231 | 5.2% | WT: BBG06471.1<br>vac2: AVW82894.1 |
| P<sub>CT</sub> | 2 | 276 | 0.7% | WT: BBG06471.1<br>vac2: AVW82894.1 |
| V<sub>CT</sub> | 1 | 68 | 1.5% | WT: BBG06472.1<br>vac2: AVW82893.1 |
| C | 5 | 186 | 2.7% | WT: BBG06473.1<br>vac2: AVW82892.1 |
| M | 5 | 335 | 1.5% | WT: BBG06474.1<br>vac2: AVW82888.1 |
| F | 3 | 550 | 0.5% | WT: BBG06475.1<br>vac2: AVW82889.1 |
| H | 17 | 617 | 2.8% | WT: BBG06476.1<br>vac2: AVW82895.1 |
| L | 29 | 2183 | 1.3% | WT: BBG06477.1<br>vac2: AVW82891.1 |
| **Total** | **101** | **5202** | **1.9%** |  |

**Table S2.**
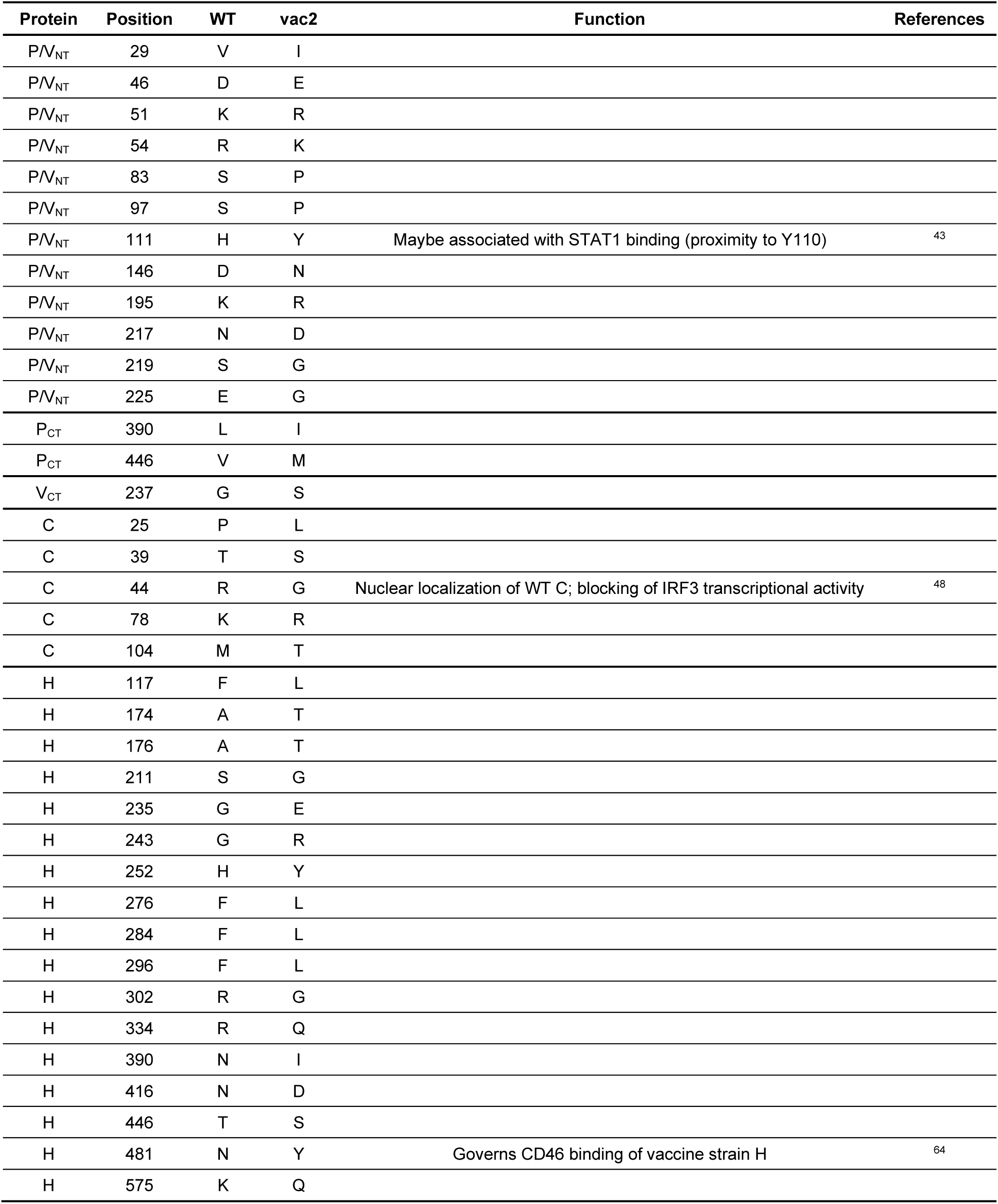
Amino acid alterations between WT and vac2 in P, V, C, and H proteins. Known functional implications are indicated. P and V proteins are divided into common amino-terminal domain (PVNT) and distinct carboxy-terminal domains (PCT, VCT).

| Protein | Position | WT | vac2 | Function | References |
| --- | --- | --- | --- | --- | --- |
| PV<sub>NT</sub> | 29 | V | I |  |  |
| PV<sub>NT</sub> | 46 | D | E |  |  |
| PV<sub>NT</sub> | 51 | K | R |  |  |
| PV<sub>NT</sub> | 54 | R | K |  |  |
| PV<sub>NT</sub> | 83 | S | P |  |  |
| PV<sub>NT</sub> | 97 | S | P |  |  |
| PV<sub>NT</sub> | 111 | H | Y | Maybe associated with STAT1 binding (proximity to Y110) | 43 |
| PV<sub>NT</sub> | 146 | D | N |  |  |
| PV<sub>NT</sub> | 195 | K | R |  |  |
| PV<sub>NT</sub> | 217 | N | D |  |  |
| PV<sub>NT</sub> | 219 | S | G |  |  |
| PV<sub>NT</sub> | 225 | E | G |  |  |
| P<sub>CT</sub> | 390 | L | I |  |  |
| P<sub>CT</sub> | 446 | V | M |  |  |
| V<sub>CT</sub> | 237 | G | S |  |  |
| C | 25 | P | L |  |  |
| C | 39 | T | S |  |  |
| C | 44 | R | G | Nuclear localization of WT C; blocking of IRF3 transcriptional activity | 48 |
| C | 78 | K | R |  |  |
| C | 104 | M | T |  |  |
| H | 117 | F | L |  |  |
| H | 174 | A | T |  |  |
| H | 176 | A | T |  |  |
| H | 211 | S | G |  |  |
| H | 235 | G | E |  |  |
| H | 243 | G | R |  |  |
| H | 252 | H | Y |  |  |
| H | 276 | F | L |  |  |
| H | 284 | F | L |  |  |
| H | 296 | F | L |  |  |
| H | 302 | R | G |  |  |
| H | 334 | R | Q |  |  |
| H | 390 | N | I |  |  |
| H | 416 | N | D |  |  |
| H | 446 | T | S |  |  |
| H | 481 | N | Y | Governs CD46 binding of vaccine strain H | 64 |
| H | 575 | K | Q |  |  |

**Table S3.**
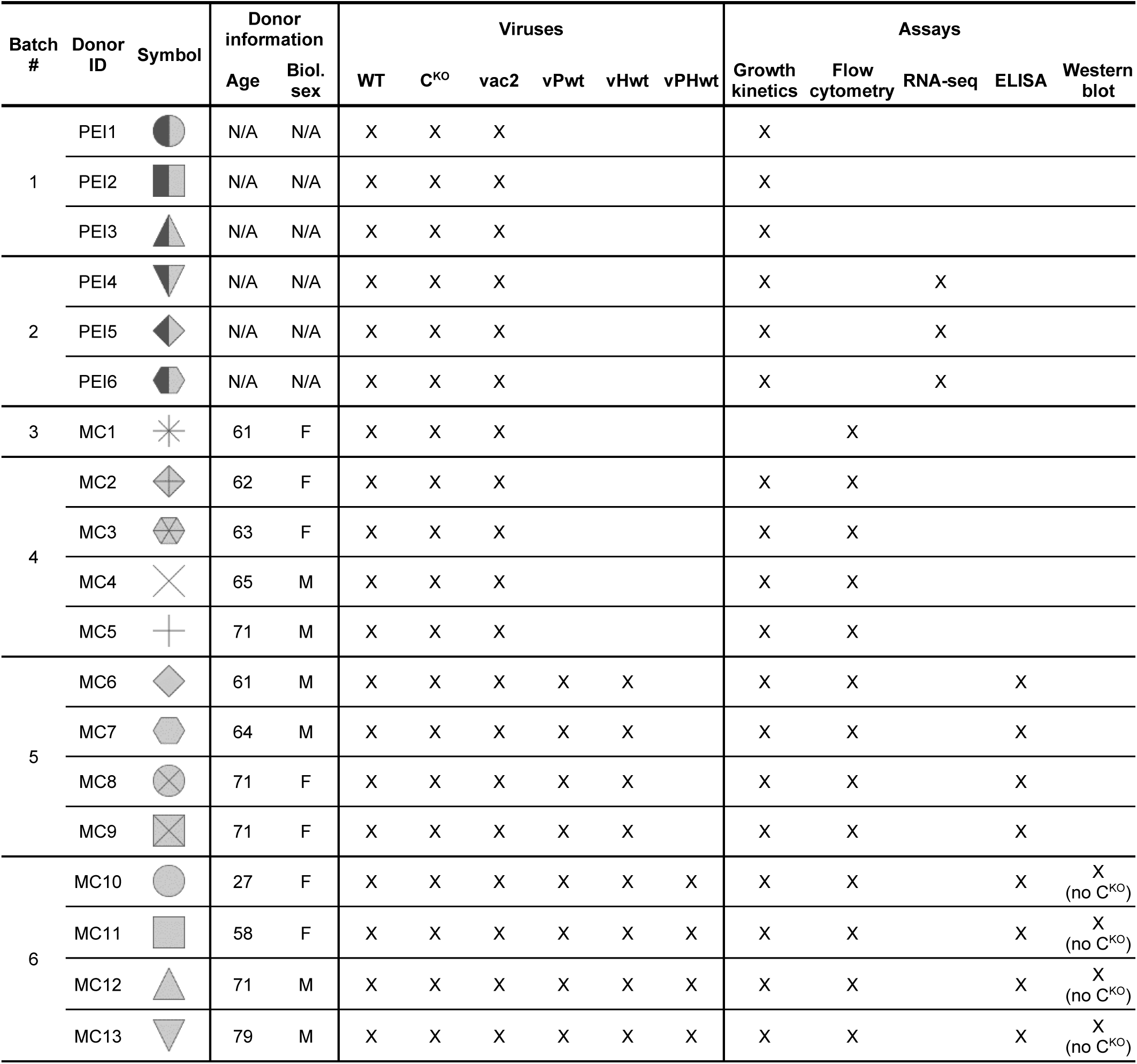
PBMC donor information summary.

| Batch # | Donor ID | Symbol | Donor information |  | Viruses |  |  |  |  |  | Assays |  |  |  |  |
| --- | --- | --- | --- | --- | --- | --- | --- | --- | --- | --- | --- | --- | --- | --- | --- |
|  |  |  | Age | Biol. sex | WT | C<sup>KO</sup> | vac2 | vPwt | vHwt | vPHwt | Growth kinetics | Flow cytometry | RNA-seq | ELISA | Western blot |
| 1 | PEI1 |  | N/A | N/A | X | X | X |  |  |  | X |  |  |  |  |
|  | PEI2 |  | N/A | N/A | X | X | X |  |  |  | X |  |  |  |  |
|  | PEI3 |  | N/A | N/A | X | X | X |  |  |  | X |  |  |  |  |
| 2 | PEI4 |  | N/A | N/A | X | X | X |  |  |  | X |  | X |  |  |
|  | PEI5 |  | N/A | N/A | X | X | X |  |  |  | X |  | X |  |  |
|  | PEI6 |  | N/A | N/A | X | X | X |  |  |  | X |  | X |  |  |
| 3 | MC1 |  | 61 | F | X | X | X |  |  |  |  | X |  |  |  |
| 4 | MC2 |  | 62 | F | X | X | X |  |  |  | X | X |  |  |  |
|  | MC3 |  | 63 | F | X | X | X |  |  |  | X | X |  |  |  |
|  | MC4 |  | 65 | M | X | X | X |  |  |  | X | X |  |  |  |
|  | MC5 |  | 71 | M | X | X | X |  |  |  | X | X |  |  |  |
| 5 | MC6 |  | 61 | M | X | X | X | X | X |  | X | X |  | X |  |
|  | MC7 |  | 64 | M | X | X | X | X | X |  | X | X |  | X |  |
|  | MC8 |  | 71 | F | X | X | X | X | X |  | X | X |  | X |  |
|  | MC9 |  | 71 | F | X | X | X | X | X |  | X | X |  | X |  |
| 6 | MC10 |  | 27 | F | X | X | X | X | X | X | X | X |  | X | X<br>(no C<sup>KO</sup>) |
|  | MC11 |  | 58 | F | X | X | X | X | X | X | X | X |  | X | X<br>(no C<sup>KO</sup>) |
|  | MC12 |  | 71 | M | X | X | X | X | X | X | X | X |  | X | X<br>(no C<sup>KO</sup>) |
|  | MC13 |  | 79 | M | X | X | X | X | X | X | X | X |  | X | X<br>(no C<sup>KO</sup>) |

